# From General-Purpose to Disease-Specific Features: Aligning LLM Embeddings on a Disease-Specific Biomedical Knowledge Graph for Drug Repurposing

**DOI:** 10.64898/2026.03.07.707871

**Authors:** Suman Pandey, Muhammed Talo, David P. Siderovski, Nathalie Sumien, Serdar Bozdag

## Abstract

Identifying new therapeutic uses for existing drugs is a major challenge in biomedicine, especially for complex neurodegenerative conditions such as Alzheimer disease and related dementias (ADRD), where treatment options remain limited and relevant data are often sparse, heterogeneous, and difficult to integrate. Although general-purpose Large Language Model (LLM) embeddings encode rich semantic information, they often lack the task-specific biomedical context needed for inference tasks such as computational drug repurposing. We introduce Contextualizing LLM Embeddings via Attention-based gRaph learning (CLEAR), a multimodal representation-fusion framework that aligns LLM embeddings with the topological structure of a context-specific Knowledge Graph (KG). Across five benchmark datasets, CLEAR achieved state-of-the-art results, improving predictive performance (e.g., F1 score) by up to 30% over prior methods. We further applied CLEAR to identify FDA-approved drugs with potential for repurposing for ADRD, including Parkinson disease-related dementia and Lewy Body dementia. CLEAR learned a biologically coherent embedding space, prioritized leading ADRD drug candidates, and accurately summarized known therapeutic relationships for FDA-approved Alzheimer disease drugs. Overall, CLEAR shows that grounding biomedical LLM embeddings with context-specific KG signals can improve drug repurposing in data-sparse, real-world settings.

GitHub: https://github.com/bozdaglab/CLEAR

## Introduction

Traditional drug discovery is a lengthy and costly process and typically takes ∼13 years and costs ∼$2–$3 billion^1,2^. For complex neurodegenerative diseases such as Alzheimer disease (AD), costs can exceed $5 billion to bring a new agent to market^3^. These long timelines and high costs, combined with stringent Food and Drug Administration (FDA) requirements for establishing efficacy and safety, can discourage investment in new therapeutics for many conditions^4^. Drug repurposing (DR) offers a viable alternative: identifying new therapeutic indications for FDA-approved drugs, enabling shorter timelines with lower cost and resource demands^5^.

Computational approaches have been critical across multiple stages of *de novo* drug discovery and of DR. They accelerate development by enabling inexpensive, rapid, large-scale virtual screening of compound libraries and drug databases, narrowing the search space^6^ and potentially identifying existing therapeutics with desired properties against new targets^7,8^. Recent advances in high-throughput molecular and clinical technologies have generated massive, publicly available biomedical and chemical datasets^9,10^. In this new era of rapidly advancing high-performance computation, driven in part by GPU-accelerated architectures optimized for large-scale data processing and model inference, one can now integrate these datasets efficiently and at scale for computational DR.

Computational DR has become a core strategy for accelerating therapy discovery, with methods aimed at predicting novel therapeutic uses for existing FDA-approved drugs^11^. Early work focused on machine learning and, matrix completion/factorization^12,13^, which infer missing entries under the assumption that similar drugs treat similar diseases. Methods such as BNNR^14^, SCMFDD^15^, and DRIMC^16^ extend this paradigm but are often limited to capturing linear associations, which may not fully represent the complex, non-linear dynamics inherent in biological systems. Network-propagation approaches then leveraged graph topology to model non-linear dynamics between drugs and diseases^17,18,19^. MBiRW^17^ uses bi-random walks to explore the local neighborhoods of drugs and diseases. iDrug^18^ learns cross-network embeddings to transfer information between related graphs, directly applying the guilt-by-association principle. DREAMwalk^19^ introduced a semantic, multi-layer guilt-by-association approach that uses guided random walks to integrate multiple levels of biological information, from molecular to semantic.

Recently, the field has shifted from tabular similarity modeling to graph-based deep learning frameworks, particularly Graph Neural Networks (GNNs). These frameworks learn complex, non-linear topological patterns more effectively^20,21^. This makes them well suited to the inherently complex structure of biomedical knowledge graphs (KGs), specifically for modeling drug-disease relationships for DR. The expressive power of GNNs makes it possible to build architectures that combine diverse biomedical information. For example, DRHGCN^20^ leveraged Graph Convolutional Networks (GCN) to capture both intra-domain (drug-drug, disease–disease) and inter-domain (drug–disease) topological relationships, enabling the model to learn rich, domain-aware embeddings that predicts novel drug–disease associations. Building on earlier GCN-based approaches, DRAGNN^21^ further enhanced representation learning by using an attention-based GNN to capture more informative relationships between drugs and diseases.

More recently, methods have begun to integrate features directly from Large Language Models (LLMs). AMVL^22^, for instance, combines LLM representations with other modalities, such as chemical-induced transcriptional profiles, to boost predictive accuracy. It utilizes embeddings from textual drug and disease descriptions where drug descriptions are obtained from DrugBank (summary, background, and chemical formula), while disease descriptions are derived from Online Mendelian Inheritance in Man (OMIM) titles and both encoded using an OpenAI embedding model^22^. Taken together, this progression reflects a clear shift from simple statistical models to expressive, multimodal deep-learning architectures that better capture biological complexity.

Despite these advances, the following key limitations persist:

a. Many computational DR methods fail to exploit the biological and pharmacological knowledge embedded in modern LLMs. Trained on large biomedical corpora, these LLMs map heterogeneous biomedical entities into a high-dimensional semantic space. This yields rich features that provide a stronger starting point for downstream modeling^23^. In contrast, many current computational DR methods either ignore such semantic features altogether or rely on representations learned through network-propagation methods (e.g., random walk-based techniques), which learn only from comparatively smaller KG (compared to large biomedical corpora) and therefore encode a much narrower scope of information.
b. Most computational DR methods rely on a narrow set of modalities and omit critical protein-level signals such as drug-protein interactions, disease-protein associations, and protein-protein interactions features. Proteins are central to drug efficacy as most drugs bind to disease etiology-associated proteins to alter pathological processes^24^. Excluding these signals could lower predictive performance in DR.
c. Most existing computational DR methods lack a robust mechanism to integrate multi-modal features, such as biomedical LLM-derived embeddings with features obtained from biomedical KGs. General-purpose LLM features often lack disease-specific context and encode biomedical entities in different, incompatible high dimension space^25^. Meanwhile, signals obtained from biomedical association/interaction networks reflect different underlying data distributions. Therefore, a robust mechanism is needed to integrate these diverse features and signals into a shared representation space, improving cross-compatibility.
d. Most computational DR methods are developed and evaluated on benchmark datasets that span many disease classes (e.g., neurodegenerative and cardiovascular diseases). In that setting, weak but crucial disease-specific signals may become diluted or are absent entirely, which could lower DR accuracy. For complex neurodegenerative disorders such as AD and related dementias (ADRD), disease-specific contextualization is essential for reliable prediction^26^.

To address the limitations of existing methods, we developed CLEAR: Contextualizing LLM Embeddings via Attention-based gRaph learning. CLEAR is a GNN-based framework that integrates and aligns general-purpose LLM embeddings/features within a disease category-specific KG. This creates a unified, context-aware feature space for downstream prediction tasks. Biomedical KG topology offers the structure needed to align incompatible high-dimensional spaces and inject disease-specific context missing from general-purpose LLM features.

In this work, as a case study, we applied CLEAR to align drug-, disease-, and protein-related LLM features with an ADRD-specific KG capturing relationships among 2,285 FDA-approved drugs, 912 neurodegenerative diseases, and 4,042 therapeutic target proteins. We then used the aligned features to predict biologically plausible candidate drugs for AD and related dementias. We demonstrate that by aligning LLM embeddings in a context-specific KG, our novel approach makes several key contributions. First, we show that CLEAR learns a biologically coherent feature space that outperforms the initial LLM features. It effectively identifies and ranks drug candidates that have biological relevance to ADRD. It further confirms known therapeutic relationships between AD and its FDA-approved AD drugs in the learned feature space. The findings of CLEAR are corroborated by published literature. Second, we establish that CLEAR achieves state-of-the-art (SOTA) performance on five benchmark drug-disease association prediction tasks, with significant gains in F1 score, a key metric for translational research.

Overall, we demonstrate an effective way to align biomedical LLM representations with KG signals to enhance tasks like computational DR. CLEAR can be applied across different disease categories (e.g., cardiovascular, autoimmune diseases, and metabolic disorders) to efficiently screen and prioritize promising treatments.

## Results

### Overview of the CLEAR framework for ADRD drug repurposing

To address the challenges of contextualizing general-purpose LLM embeddings, we developed CLEAR: **C**ontextualizing **L**LM **E**mbeddings via **A**ttention-based g**R**aph learning. CLEAR is a framework that integrates and aligns general-purpose LLM embeddings within a disease category-specific KG to identify and rank novel drug-disease associations. In this study, we built a neurogenerative disease-specific KD. As illustrated in **Fig. 1**, the CLEAR framework operates as a sequential pipeline comprised of five modules.

1. An attributed ADRD KG is constructed, integrating FDA-approved drugs, neurodegenerative diseases, and their therapeutic target proteins associated with these drugs and diseases as nodes in the graph, with edges representing known associations and similarities curated from public databases as shown in **Table 1** (**Fig. 1a**).
2. Each node is initialized with a rich feature vector generated from pretrained LLMs: drug nodes with Simplified Molecular Input Line Entry System (SMILES)-based features from MoLFormer^28^, disease nodes with descriptive features from BioBERT^29^, and protein nodes with UniProt protein sequence-based features from ESM-2^30^ (**Fig. 1b**).
3. CLEAR refines these initial features by learning ADRD KG’s topology. It applies multi-relational Graph Attention Networks (GATs) to generate context-specific feature updates for each node based on its different relationship types (e.g., drug-disease, drug-protein, and drug-drug similarity). A Multi-Head Self-Attention (MHSA) mechanism then learns to fuse these multiple, topologically informed features into a single, unified embedding for each node, which we term as CLEAR embedding (**Fig. 1c**).
4. These CLEAR embeddings are used to train a two-layer multi-layer perceptron (MLP) for link prediction, which performs KG completion task by inferring novel, high-confidence bipartite links (i.e., drug-disease, drug-protein, and disease-protein). This model is trained on known associations as positive examples and challenging negatives generated via a topology-aware sampling (**Fig. 1d**).
5. The completed KG and CLEAR embeddings serve as the basis for downstream applications, such as generating a prioritized list of DR candidates for a user-queried disease within in KG via a network topology-based ranking algorithm (**Fig. 1e**). In this study, we queried three ADRDs to create a prioritized list of DR candidates.

**Fig. 1.**
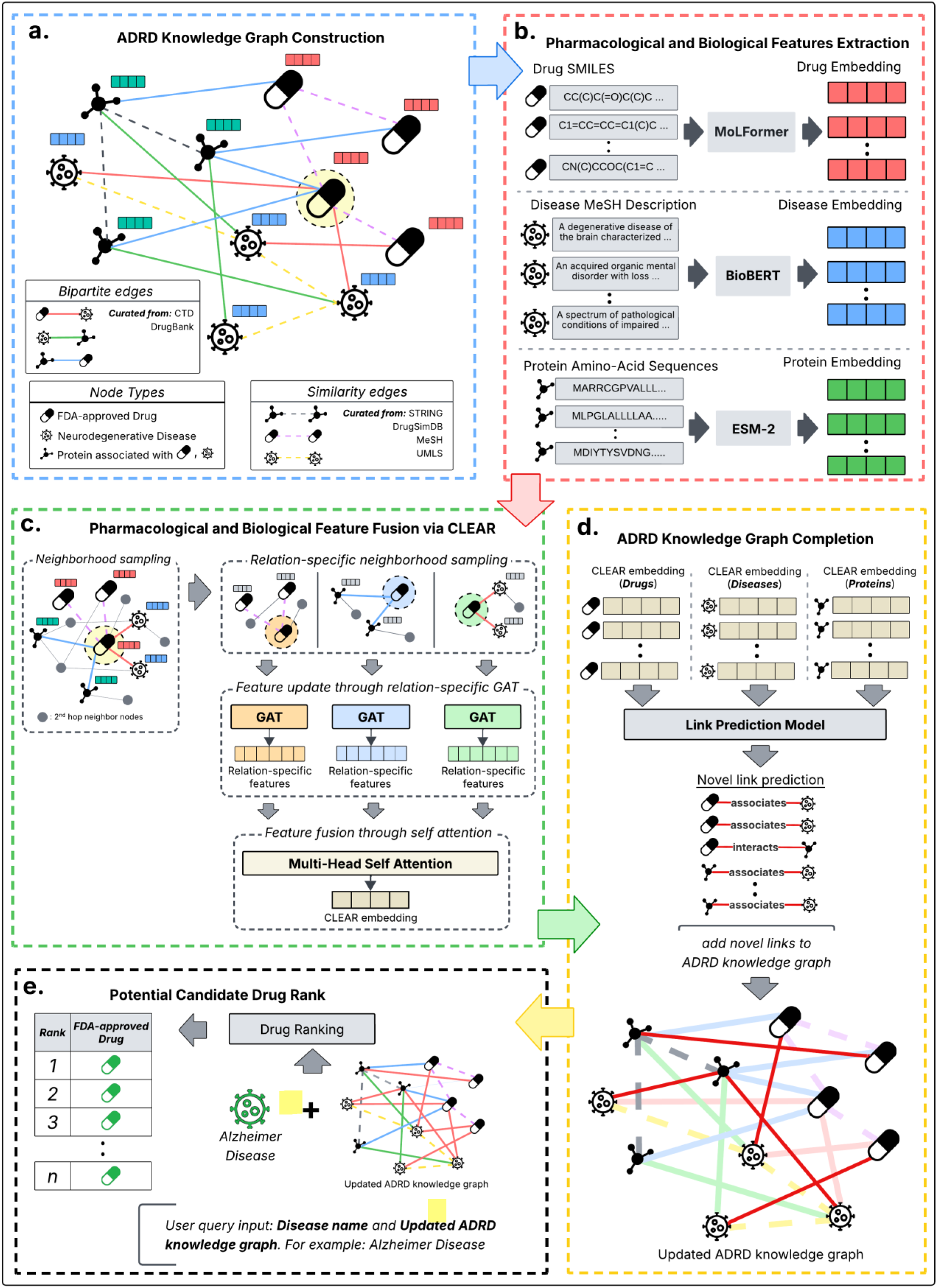
Overview of the CLEAR framework. **a.** First, the Alzheimer Disease and Related Dementias (ADRD) KG is constructed. The KG composed of nodes representing FDA-approved drugs, neurodegenerative diseases, and associated proteins, and the edges representing known associations between nodes curated from public databases. **b**. Each node in this graph is then initialized with rich features extracted using pretrained large language models (LLMs): Drug nodes are initialized with SMILES-based features, disease nodes are initialized with disease description-based features, and protein nodes are initialized with protein sequence-based features. **c.** Operating on this attributed graph, CLEAR learns to update LLM-based node features through generating multiple relation-specific topology feature updates followed by multi-relation feature fusion. **d.** The updated features are then used to train a link predictor, which performs KG completion by inferring novel, high-confidence bipartite links (i.e., drug-disease, drug-protein, and disease-protein links). **e.** Finally, this completed and enriched graph serves as the basis for downstream applications, allowing a user to query a specific disease and generate a prioritized list of potential drug candidates.

**Table 1.** Composition of the ADRD KG. The number of associations and corresponding data sources are listed for all six relation types, comprising both similarity and bipartite links.

| Relation types | Number of associations | Data source |
| --- | --- | --- |
| Drug-disease association | 11,354 | CTD |
| Disease-protein association | 3,403 | DrugBank |
| Drug-protein association | 11,818 | CTD |
| Drug-drug similarity | 75,704 | DrugSimDB |
| Disease-disease similarity | 183,828 | MeSH, UMLS |
| Protein-protein similarity | 74,556 | STRING |

#FDA-approved **Drug** node: 2,285
#Neurodegenerative **Disease** nodes: 912
#Therapeutic target **Protein** nodes: 4,042 (#Unique therapeutic target proteins of FDA-approved drugs: 2,265, #Unique therapeutic target proteins associated with neurodegenerative diseases: 1,147, #Common therapeutic target proteins: 630)

### CLEAR achieves SOTA performance across multiple benchmark datasets and tasks

To evaluate our framework, we benchmarked CLEAR’s performance on three fronts:

First, we assessed CLEAR’s capacity to learn from the entire, multi-relational ADRD KG. In a ten-fold cross-validation setup, CLEAR was tasked with predicting all six types of relationships, including drug-disease, drug-protein, disease-protein, and all corresponding similarity links. As shown in **Supplementary Figure 1**, the model demonstrated robust performance across all tasks based on the commonly used evaluation metrics, achieving an average F1 score over 0.85, and Area Under Precision Recall Curve (AUPR) and Area Under Receiver Operating Characteristics (AUCROC) scores over 0.93. The range of these metrics is 0-1. For more information about them see **Supplementary Notes 1**. These results indicate that CLEAR effectively learns a unified embedding space that captures the complex interplay between all entities in the graph.

Second, to evaluate the generalizability of our framework, we tested it on five widely used drug-disease prediction benchmark datasets, namely Cdataset^17^, Fdataset^31^, Ydataset^32^, LAGCN^33^, and LRSSL^34^. The statistics of these benchmark datasets are described in **Supplementary Table 1.**

These datasets are all derived from different snapshots of the Comparative Toxicogenomic Database (CTD)^35^, making them suitable for assessing model robustness to data distribution shifts. As illustrated in **Supplementary Figure 2**, these benchmark datasets have minimal overlaps (∼10% on average) with each other and with ADRD KG’s drug-disease prediction task, confirming they represent distinct prediction tasks. For this analysis, we compared CLEAR against the top five performing SOTA methods for each specific dataset, as identified in a comprehensive review paper^36^. The SOTA methods used for comparison on each dataset were as follows:

- Cdataset: ITRPCA^37^, MLMC^38^, OMC^39^, VDA-GKSBMF^40^, BNNR
- Fdataset: HINGRL^41^, ITRPCA, MLMC, OMC, DRRS^42^
- Ydataset: VDA-GKSBMF, DRRS, ITRPCA, DRPADC^43^, HGIMC^44^
- LAGCN: SCMFDD, GROBMC^45^, HINGRL, LAGCN, DRHGCN
- LRSSL: GROBMC, DRHGCN, DRIMC, OMC, DDAPRED^46^

The same data curation and filtration protocol used for the ADRD KG was applied to each benchmark dataset. A persistent challenge with existing SOTA methods is that their performance has largely stagnated; while many achieve high AUCROC scores (often >0.95), their F1 scores are typically much lower^36^. This indicates a high false-positive rate when a decision threshold is applied, which is a critical flaw for computational DR, where the goal is to reliably filter candidates before conducting expensive experimental validation. In contrast, CLEAR demonstrated the best performance, particularly in the F1 score, without any dataset-specific hyperparameter tuning. It achieved an F1 score improvement of ∼26% on Cdataset, ∼8% on Fdataset, ∼23% on Ydataset, ∼2% on LAGCN, and ∼30% on LRSSL over the next-best SOTA method (**Fig. 2c**–**2g, Supplementary Table 2-6**). Even though the AUCROC and AUPR scores of the SOTA methods were saturated (i.e., >0.95 on average), CLEAR achieved improvements above this saturated state (i.e., ∼1-2% improvements across all datasets tested). This superior performance, achieved with the same set of hyperparameters across datasets, underscores that CLEAR is a more robust and generalizable framework. To further assess its generalizability, we evaluated CLEAR’s ability to model all six relationship types simultaneously on the benchmark datasets. On this multi-task evaluation, CLEAR consistently achieved an F1 score greater than 0.80 (**Supplementary Figure 1**) for all relationship types across all datasets. This result demonstrates CLEAR’s capacity to generalize effectively not only to new datasets but also to diverse prediction tasks.

**Fig. 2.**
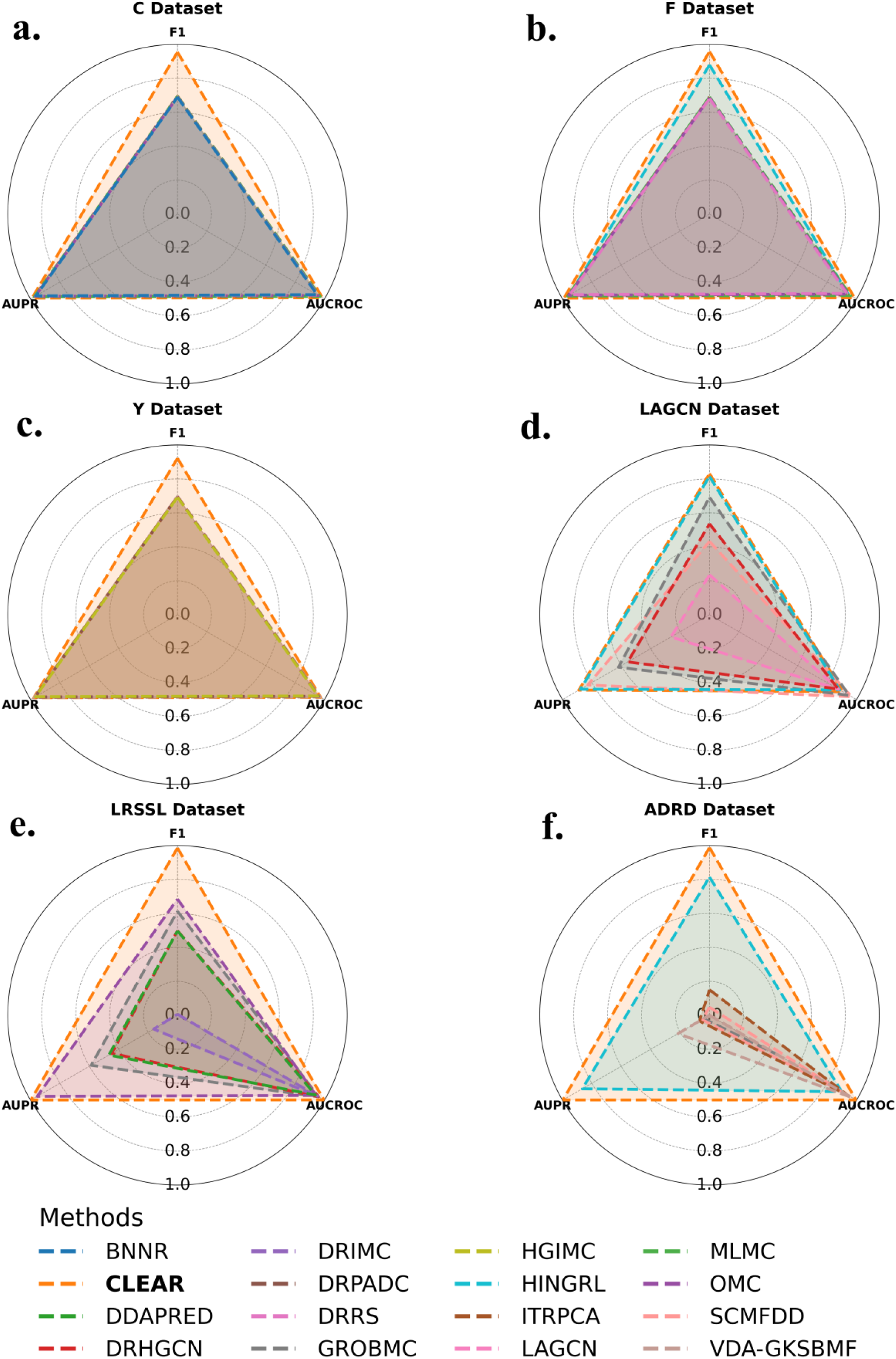
Comparative performance analysis of CLEAR across multiple datasets. Model performance is visualized using radar plots, where the three axes represent F1, AUPR, and AUCROC scores and points that are further from the center indicate higher performance score. **a. - e.** Comparison of CLEAR against the top SOTA methods on five benchmark datasets, **a.** Cdataset, **b.** Fdataset, **c.** Ydataset, **d.** LAGCN, and **e.** LRSSL. **f.** Comparison of CLEAR’s performance against SOTA methods on the drug-disease association prediction task of the ADRD dataset. All results represent the average performance from a 10-fold cross-validation.

Finally, to evaluate the SOTA method’s performance on predicting novel drug-disease relationships from the ADRD KG, we picked the SOTA method for each of the five benchmark datasets as identified in a comprehensive review paper^36^ (**Fig. 2b**). Among these methods, HINGRL achieved the second-best performance with an F1 score of 0.815 and an AUCROC of 0.899 (**Supplementary Table 7**). However, most other methods, while attaining high AUCROC scores, yielded low F1 and AUPR scores (often <0.2). CLEAR significantly outperformed all of them, with an F1 score of 0.989 and an AUCROC of 0.996. This performance gap suggests that, for a specialized and complex task like ADRD-focused computational DR, a model’s ability to integrate rich, multi-modal information (especially protein-related data and context-aware embeddings) is crucial.

### CLEAR identifies novel candidate drugs for ADRD

To evaluate our framework, we examined CLEAR’s ability to identify biologically plausible novel drug candidates for ADRD while learning a feature space that accurately captures known pharmacological and biological relationships. Specifically, we applied CLEAR to prioritize potential drug candidates for ADRD such as AD, Parkinson Disease Dementia (PDD), and Lewy Body Dementia (LBD). CLEAR effectively ranks drug candidates that, in *post hoc* analyses employing human-curated knowledge of published biomedical literature, are seen to have *bona fide* biological relevance to these diseases (**Table 2**).

**Table 2.** Top five drug repurposing candidates for ADRD: Alzheimer Disease, Parkinson Disease Dementia, and Lewy Body Dementia. The table lists the top five candidate drugs predicted by CLEAR, their known biological properties relevant to ADRD, and *post hoc*-identified evidence from the biomedical literature supporting their potential clinical association with ADRD.

|  | Rank | Drug (substance)<br>name | *Post hoc*<br>Justification | Literature<br>Citations |
| --- | --- | --- | --- | --- |
| | 1 | Zinc | Zinc dyshomeostasis promotes Aβ aggregation and oxidative stress in AD | 47,48,49 |
| *Alzheimer Disease* | 2 | Copper | Copper imbalance linked to amyloid toxicity and oxidative neuronal injury | 50,51 |
|  | 3 | Dextromethorphan | NMDA/sigma-1 actions confer neuroprotection; DXM use for other indications reported to lower dementia risk | 52,53 |
| | 4 | Melatonin | Antioxidant: reduces Aβ toxicity and supports circadian-linked neuroprotection | 54,55,56 |
|  | 5 | Physostigmine | Prototype cholinesterase inhibitor; precursor to current AD cognitive therapies | 57 |
| *Parkinson Disease Dementia* | 1 | Dopamine | Dopaminergic replacement remains cornerstone of PD and PDD management | 58,59 |
| | 2 | Copper | Copper dysregulation promotes α-synuclein aggregation and oxidative damage | 60,61 |
|  | 3 | Glutathione | Early PD marker: antioxidant depletion linked to nigral neuron loss | 62,63,64 |
|  | 4 | Zinc | Zinc affects dopaminergic neuronal survival and redox homeostasis in PD | 65,66,67 |
|  | 5 | Loxapine | Serotonin/dopamine modulator used for psychosis in PD spectrum | 68 |
| *Lewy Body Dementia* | 1 | Amitriptyline | Used for mood/sleep symptoms; cholinergic burden limits use | 69 |
| | 2 | Estradiol | Estrogen signaling shown to mitigate α-synuclein toxicity | 70 |
| | 3 | Zinc | Metal imbalance contributes to α-synuclein aggregation in LBD | 71 |
|  | 4 | Paclitaxel | Microtubule stabilization may counter axonal transport deficits | 72 |
| | 5 | Clenbuterol | β-adrenergic agonist with neurotrophic and anti-inflammatory actions | 73 |

To determine whether CLEAR’s top-ranked drugs share mechanistic links to the disease, we further considered one of the top-ranked novel drug candidates for AD: Dextromethorphan. Dextromethorphan (DXM), a long-used antitussive agent, has a multifaceted CNS pharmacology. It acts as a noncompetitive NMDA receptor antagonist, a property that has been linked to modulation of excitotoxic glutamatergic signaling and neuroprotection in preclinical studies^74,75^ and is shared with Memantine (NMDA receptor antagonist approved for moderate-to-severe AD)^76,77^. DXM is also a sigma-1 receptor agonist^78^, which has been implicated in neuromodulatory and putative neuroprotective effects^79,80^.

DXM is also involved in the modulation of monoaminergic neurotransmission, notably through clinical evidence of its inhibition of serotonin reuptake ^81,82^, overlapping mechanistically with the serotonin-dopamine activity of Brexpiprazole (FDA approved for AD-associated agitation)^83,84^. Moreover, DXM and its primary metabolite dextrorphan are seen to block nicotinic acetylcholine receptor function, particularly at α3β4-containing subtypes, in both *in vitro* and *in vivo* experimental systems^85,86^. These mechanisms contribute to its complex neuropharmacologic profile and are relevant when considering DXM’s broader CNS effects beyond cough suppression; for example, a fixed-dose combination of DXM and bupropion was FDA-approved in 2022 for the treatment of major depressive disorder (MDD) in adults, reflecting rapid improvements in depressive symptoms compared with placebo in phase 3 trials of patients with MDD^87^.

Although not currently approved for AD, DXM’s biological relevance is supported by CLEAR-generated overlap in its therapeutic targets with proteins implicated in AD pathology. The ADRD KG contains 101 proteins associated with AD and 21 proteins targeted by DXM. Of these, three proteins are shared between the protein-disease and protein-drug association classes: neuronal acetylcholine receptor subunit alpha-7 (UniProt P36544, encoded by *CHRNA7*), neuronal acetylcholine receptor subunit beta-2 (UniProt P17787, encoded by *CHRNB2*), and membrane-associated progesterone receptor component 1 (UniProt O00264, encoded by *PGRMC1*). The involvement of these specific proteins in AD is well-documented. Neuronal nicotinic acetylcholine receptors (nAChRs), including those containing α7 or β2 subunits, represent major therapeutic targets for restoring cholinergic tone and cognitive function in AD, consistent with the mechanisms of the approved acetylcholinesterase inhibitors donepezil, rivastigmine, and galantamine, the latter also acting directly as a positive allosteric modulator of α7-containing nAChRs)^88,89,90^. Membrane-associated progesterone receptor component 1 (PGRMC1) participates in amyloid-β–related pathology through its role in a multiprotein complex that mediates Aβ₄₂ binding and neuronal uptake; early work suggested that PGRMC1 itself functioned as the sigma-2 receptor mediating Aβ oligomer toxicity^91^, but subsequent studies demonstrated that the true sigma-2 receptor is TMEM97, which forms a complex with PGRMC1 and the LDL receptor to internalize Aβ₄₂ and its aggregates^92^. The purported mechanistic link between Aβ binding and neuronal toxicity continues to drive the translational focus on Aβ-directed therapeutics, exemplified by recent FDA approvals of monoclonal antibodies such as aducanumab, lecanemab, and donanemab, which target aggregated or protofibrillar Aβ species to reduce amyloid burden in patients with early AD^93,94,95^.

Further supporting the value of CLEAR’s outcomes, Gene Ontology (GO) enrichment analysis of Dextromethorphan’s 21 protein targets reveals a significant over-representation of terms highly relevant to AD (**Supplementary Table 8-10**). The most enriched biological process is “respiratory burst (GO:0045730),” a key component of the microglial activation that drives chronic neuroinflammation in AD^96,97^. Similarly, “cholinergic synaptic transmission (GO:0007271)” is a critical process, as the degeneration of cholinergic neurons is a hallmark of AD that leads to cognitive decline^98,99^. Among molecular functions, “acetylcholine-gated cation-selective channel activity (GO:0022848)” is a top hit, directly implicating the cholinergic system that is the target of several existing AD drugs as previously mentioned. These results strongly suggest that CLEAR identified Dextromethorphan based on sound biological reasoning encoded in the KG. Notably, an independent, population-based cohort study from Taiwan ^100^ reported that DXM use for other indications was associated with a significantly lower risk of developing dementia, providing real-world epidemiologic support that complements the CLEAR-derived mechanistic prediction.

*Post-hoc* review of the biomedical literature and clinical trial registries reveals that DXM-based compounds have indeed been evaluated in clinical studies targeting agitation associated with AD, although none have attained FDA approval for this indication. A randomized, double-blind, placebo-controlled phase 2 trial of Dextromethorphan-Quinidine (compound used to stabilize DXM) demonstrated a statistically significant reduction in agitation severity among individuals with probable AD, with acceptable tolerability over 10 weeks of treatment^101^. Further efforts explored deuterated analogs of DXM combinations (e.g., AVP-786) and formulations such as AXS-05 in similar neuropsychiatric symptom domains, with mixed or ongoing results reported in literature and ClinicalTrials.gov entries for agitation protocols (e.g., NCT01584440; NCT04797715). Notably, these investigations have focused on neuropsychiatric and behavioral symptoms rather than on cognitive decline or disease modification *per se*; in contrast, the CLEAR prediction of Dextromethorphan for AD emerges from its inferred associations with core AD-relevant molecular pathways and targets implicated in cognition and neurodegeneration, suggesting that the ML-driven link is mechanistically distinct from the agitation symptom management context that has dominated clinical research into DXM’s utility to date.

### CLEAR captures known therapeutic relationships among FDA-approved drugs for AD and improves baseline classifier performance

We next assessed whether CLEAR’s embeddings represent known therapeutic relationships more effectively than the general-purpose LLM features (i.e., the initial drug, protein, and disease features learned from pretrained LLMs before applying CLEAR). In principle, therapeutically related entities such as drugs, diseases, and their associated proteins should be closer in the embedding space, as similar drugs often treat related diseases and share common mechanisms of action. To assess this, we computed the Euclidean distances between embedding vectors for all cross-pairs between the drug-side nodes (i.e., drugs, drug-target proteins) and the disease-side nodes (diseases, disease-associated proteins): drug–disease, drug–disease-associated protein, drug-target protein–disease, and drug-target protein–disease-associated protein. Specifically, we analyzed the Euclidean distances between CLEAR embeddings of five FDA-approved AD drugs (Donepezil, Galantamine, Rivastigmine, Memantine, and Brexpiprazole)^24^ and their 57 targets, and the AD node and 101 AD-associated proteins. For the general-purpose LLM feature comparison, Principal Component Analysis (PCA) was used to reduce the dimensionality of protein features to match that of drugs and diseases. The results, visualized in **Fig. 3a-d**, show that CLEAR learns a more contextualized embedding space.

**Fig. 3.**
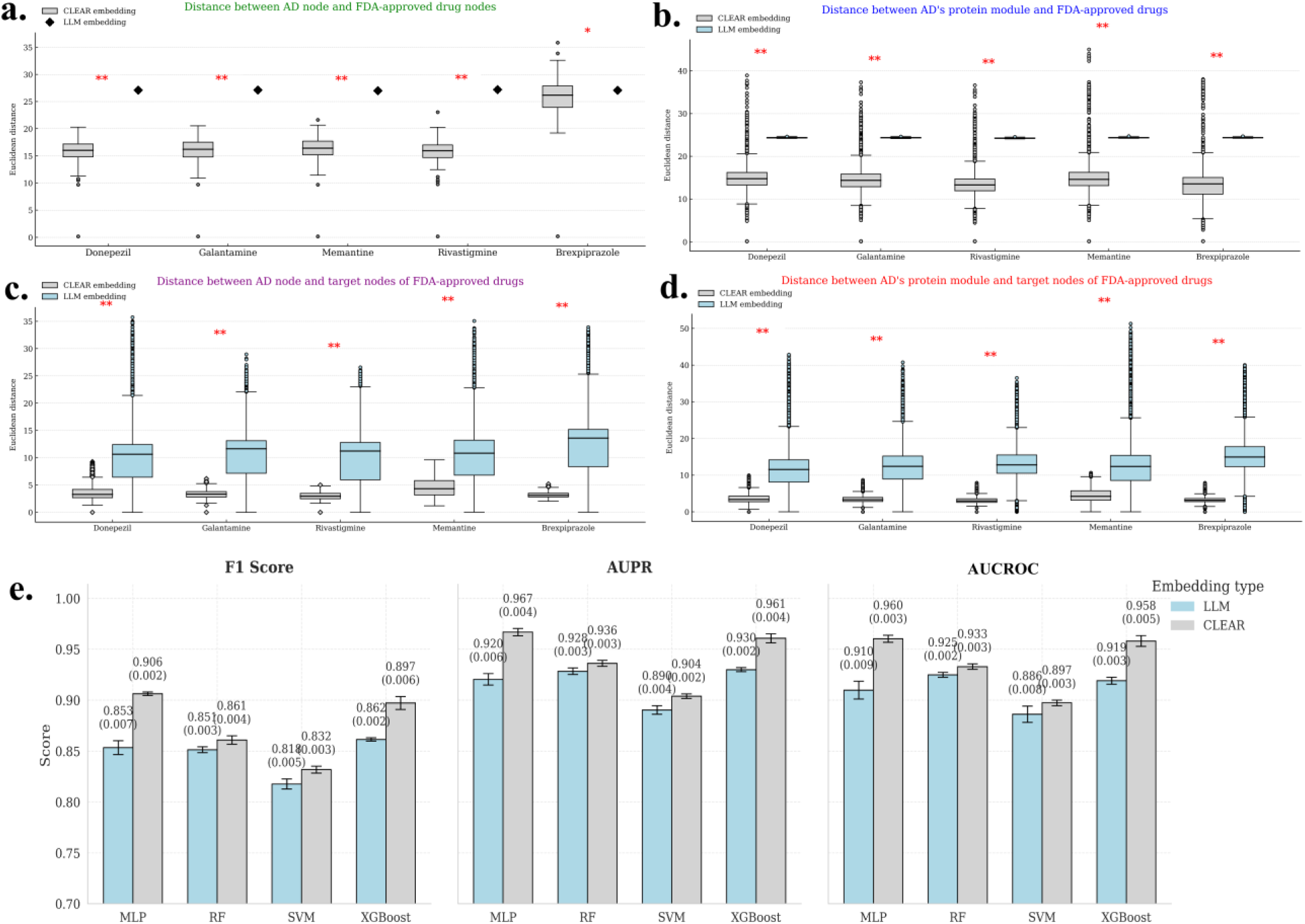
CLEAR aligns feature space to validate relationships of five FDA-approved AD drugs: Donepezil, Galantamine, Rivastigmine, Memantine, and Brexpiprazole. The distribution of distances for these five approved drugs on the general-purpose LLM embeddings versus the CLEAR embeddings space. The distances shown are, between **a.** the AD node and the FDA-approved AD drug nodes, **b.** the FDA-approved AD drug nodes and AD-associated protein nodes, **c.** the AD node and target protein nodes of FDA-approved AD drugs, **d.** target protein nodes of FDA-approved AD drugs and the AD node. *: p-value < 0.05, **: p<0.0001. **e.** Embedding performance and ablation analysis: performance comparison of machine learning classifiers (MLP, Random Forest, SVM, and XGBoost) on the ADRD drug-disease prediction task, trained using either the initial LLM- or CLEAR-embeddings. The differences in F1, AUPR, and AUROC between machine learning classifiers using LLM and CLEAR embeddings are statistically significant (F1, p = 3.35E-10; AUPR, p = 4.54E-10; AUCROC, p = 7.13E-10).

The distances between the AD and the five FDA-approved drugs are significantly smaller in the CLEAR embedding space compared to the general-purpose LLM feature space (**Fig. 3a**) (*p* ≤ 0.0001). The drugs are brought significantly closer to AD’s target module in the CLEAR embedding space (**Fig. 3b**) (*p* ≤ 0.0001). We also observed that the AD node is positioned significantly closer to all known therapeutic targets of these drugs (**Fig. 3c)** (*p* ≤ 0.0001). Finally, the target protein modules of these drugs move significantly closer to AD-related proteins in the CLEAR embedding space (**Fig. 3d)** (*p* ≤ 0.0001). These results indicate that CLEAR maintains close relationships between drugs and disease with respect to their therapeutic targets. This demonstrates its ability to integrate pharmacological information (drug target proteins) and biological information (disease-associated proteins) into a unified, context-specific representation. The general-purpose LLM features fail to capture these specific drug-disease and drug-target proteins relationships, leaving them far apart in the feature space.

Next, to establish that the embeddings produced by CLEAR are more informative and context specific for downstream tasks than the general-purpose LLM features, we trained and tested four standard ML classifiers (Random Forest (RF), SVM, MLP, and XGBoost) on the ADRD drug-disease prediction task. We trained two models for each ML classifier: one using concatenated general-purpose LLM features and the other using concatenated CLEAR embeddings for drug and disease nodes. As shown in **Fig. 3e** and **Supplementary Table 11**, using CLEAR embeddings consistently improved the performance of all four classifiers across all metrics. We observed an average performance significantly increased across all metrics (i.e., F1, AUCROC, and AUPR) of approximately 5.6% for MLP, 3.7% for XGBoost, 1.3% for SVM, and 0.9% for RF across 10-fold cross validation (F1, p = 3.35E-10; AUPR, p = 4.54E-10; AUCROC, p = 7.13E-10). This demonstrates that the biological and pharmacological information integrated by CLEAR makes the general-purpose LLM feature representation more context-specific, which in turn increases ML model performance.

### Ablation study shows CLEAR’s components are critical to its performance

We conducted a comprehensive ablation study to evaluate the importance of CLEAR’s core components. We systematically removed individual modules, both computational and data-related, to create six distinct variants of CLEAR and compared their performance against the complete CLEAR framework (**Fig. 4, Supplementary Table 12**) across 10-fold cross validation.

**Fig. 4.**
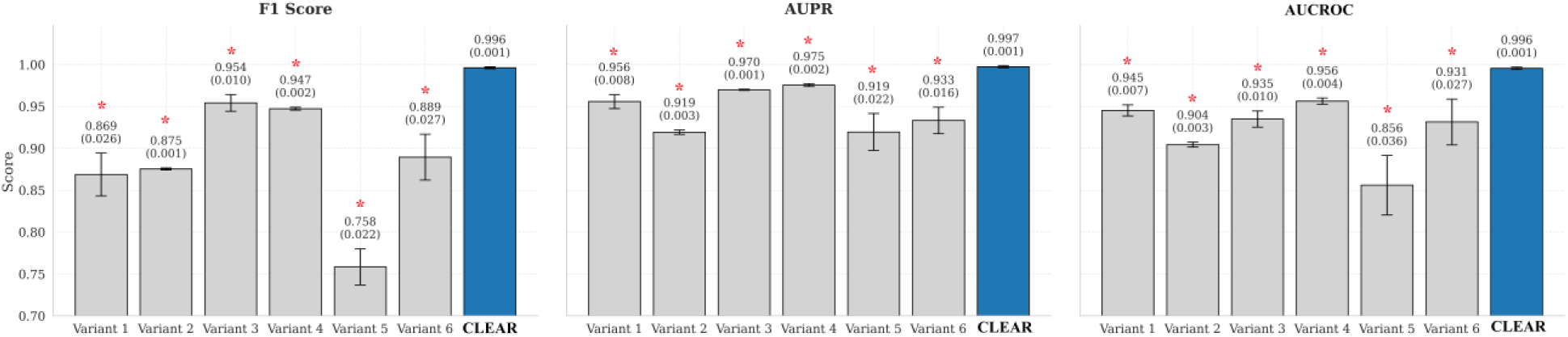
Ablation study evaluating the contribution of key components to the performance of CLEAR. CLEAR (Complete) is compared against six variants trained: without GAT and MHSA (Variant 1), without MHSA (Variant 2), using a standard, non-weighted loss function (Variant 3), using random negative sampling (Variant 4), with randomly initialized node features instead of LLM-based features (Variant 5), and using only drug- and disease-related data, excluding protein-related information (Variant 6). All results are reported as the mean (± standard deviation) from a 10-fold cross-validation. The differences in F1, AUPR, and AUCROC between the variants are statistically significant (F1, *p* ≤ 0.0001; AUPR, *p* ≤ 0.0001; AUCROC, *p* ≤ 0.0001)

The computational modules ablated were:

1. No GATs and No MHSA (Variant 1): The entire graph learning and fusion architecture were replaced with a simple 2-layer MLP trained on the concatenated general-purpose LLM features.
2. No MHSA (Variant 2): The attention-based fusion was replaced with a simple averaging of relation-specific embeddings.
3. No Weighted Loss Function (Variant 3): The weighted binary cross-entropy was replaced with a standard binary cross-entropy loss.
4. No Topology-Aware Negative Sampling (Variant 4): CLEAR’s topology-aware negative sampling was replaced with random sampling of non-existent edges in the ADRD KG.

The data modules ablated were:

1. No general-purpose LLM features (Variant 5): The general-purpose LLM features from the pretrained models were replaced with randomly initialized vectors sampled from a Gaussian distribution with a mean of 0 and a standard deviation of 1.
2. No protein Information (Variant 6): All protein nodes and their associated edges were removed from the KG.

The results demonstrate that each CLEAR’s component plays a crucial role in the final model performance (**Fig. 4**). The largest performance decrease was observed in Variant 5, where replacing the general-purpose LLM features with random features caused the F1 score to drop by ∼24%. This suggests that the general-purpose LLM features provide an essential foundation for CLEAR. The next largest performance decreases were seen in Variant 1 (no graph learning), Variant 2 (no attention-based fusion), and Variant 6 (no protein data). This highlights the importance of using protein-related information and attention-based mechanisms to extract and fuse information from multi-relationships. Finally, while the impact of removing the weighted loss (Variant 3) and topology-aware sampling (Variant 4) was less severe, their replacement still resulted in a noticeable drop in performance. The differences in F1, AUPR, and AUROC between the CLEAR (Complete) and all its variants were statistically significant (F1, *p* ≤ 0.0001; AUPR, *p* ≤ 0.0001; AUCROC, *p* ≤ 0.0001), confirming that each component contributes meaningfully to CLEAR’s overall performance. This suggests that careful management of class imbalance and the generation of challenging negative samples are important for robust training.

### CLEAR preserves drug-disease prediction performance under sparse supervision but reduces candidate drug ranking specificity

Next, we investigated how CLEAR performs when trained with sparse data. CLEAR integrates three data modalities: (a) LLM-derived node embeddings, (b) drug-disease links, and (c) other links such as disease-protein, drug-protein, and similarity links. To understand their individual and combined effects, we systematically reduced the amount of available training information across all data modalities. Specifically, we varied the proportion of drug-disease links used for training as 0%, 1%, 5%, 10%, and 25%; the proportion of other links as 0%, 1%, 5%, and 10%; and the proportion of LLM-derived node embeddings as 0%, 10%, 20%, 30%, 40%, 50%, 60%, 70%, 80%, 90%, and 100%. For each combination, only the specified fraction of each modality was used to train CLEAR. All models were trained following the same protocol as the *full CLEAR* (i.e., CLEAR trained with 100% of the training data).

The results summarized in **Supplementary Figure 3** reflect CLEAR’s performance on predicting drug-disease links in terms of F1 score, AUROC, and AUPR when trained with different combinations of sparse data modalities over 10-fold cross-validation. We observed that as expected the most critical relation type to predict drug-disease links was drug-disease. Using only 25% of the drug-disease links was sufficient for CLEAR to achieve the full CLEAR’s drug-disease prediction performance even in the cases of limited links or LLM embeddings. We also observed that LLM embeddings contributed when there were very limited drug-disease links and other links. For instance, when drug-disease links are only 1% and other links are 0% (or vice versa), using 100% of LLM embeddings increased the F1 score over 10%.

To further assess how CLEAR performs in prioritizing biologically plausible drug candidates in sparse network settings, we focused on two extreme data modality configurations: (a) training with 0% LLM-derived node embeddings and (b) with 100% LLM-derived node embeddings, while varying the proportion of other links and fixing drug-disease links at 25%. The downstream drug rankings produced under these sparse training settings remain largely unchanged across AD, PDD, and LBD suggesting that under sparse network settings drug prioritizations are not disease-specific (**Supplementary Table 13-14**). For example, as shown in **Supplementary Table 13**, Viomycin, Rituximab, and Casirivimab are consistently ranked among the top candidates for AD, PDD, and LBD. These drugs have no established neuroprotective relevance and instead are antimicrobial or immunologic therapies that are not designed to target central nervous system pathology. Furthermore, several drugs prioritized under these sparse settings belong to medication classes that prior clinical studies have shown can worsen cognitive or neuropsychiatric symptoms in dementia. As shown in **Supplementary Table 14**, Dequalinium and Diphemanil appears among top ranked drugs for AD. However, both drugs are anticholinergic agents that directly oppose cholinergic signaling deficits central to AD pathology^102,103,104^.

These observations reinforce the idea that while sparse training is sufficient to achieve strong performance on the drug-disease link prediction task, high predictive metrics do not guarantee success on downstream applications such as drug repurposing ranking. Biologically and clinically meaningful drug prioritization requires richer supervision to avoid elevating drugs that are ineffective or potentially harmful for ADRD. We also note that evaluating the correctness of downstream drug rankings is inherently subjective, as comprehensive biological or clinical validation is not feasible within the scope of this study. Consequently, our assessment of ranking quality is based on evidence from published literature and established clinical knowledge, rather than direct experimental validation.

## Discussion

LLMs produce general-purpose text-derived embeddings of biomedical entities, but these representations lack disease-specific context and reside in incompatible embedding spaces. In this study, we introduce the CLEAR framework, which implements a multi-relational GNN to contextualize and align general-purpose LLM embeddings of drugs, diseases, and protein within a biomedical KG. This creates a unified, context-aware feature space for downstream prediction tasks such as prioritizing top candidate drugs. CLEAR generates a unified, disease-specific embeddings, which we call CLEAR embeddings, by integrating multi-modal data and leveraging attention-based learning methods. Our results show that CLEAR embeddings are more informative at multiple downstream link prediction tasks compared to the general-purpose LLM embeddings.

CLEAR demonstrated high performance on the ADRD KG across various link prediction tasks (e.g., drug-disease association, drug-protein interaction, disease-protein association, and drug-similarity). Furthermore, it achieved SOTA performance on five external benchmark datasets without any dataset-specific hyperparameter tuning (**Fig. 2**). Together, these results demonstrate the generalizability and robustness of our approach. Our ablation studies confirmed that all proposed data and computational modules were essential for CLEAR’s performance. This includes the initial LLM features, the multi-relational GNN fusion, and the inclusion of protein-related information.

The case study on CLEAR’s top-ranked candidate drug for AD (Dextromethorphan), illustrates that the model’s predictions are biologically grounded, with the drug’s known therapeutic target proteins being highly relevant to AD pathology (**Supplementary Table 8-10**). Furthermore, the distance analysis of FDA-approved AD drugs demonstrated that CLEAR learns a more compact and biologically coherent embeddings compared to general-purpose LLM embedding spaces. CLEAR correctly puts therapeutically similar entities such as disease target module and drug’s therapeutic target module closer together (**Fig. 3a-d**).

However, the current CLEAR architecture, while effective, is memory intensive. This arises from the use of multiple attention-based feature update modules like relation-specific GATs and a MHSA module (**Supplementary Fig. 3**). However, as CLEAR is a GNN-based framework, sub-graph sampling could be used to scale it to larger KG or to train it on systems with limited GPU memory.

While we have used CLEAR embeddings for link prediction tasks in this study, CLEAR could be used for other downstream tasks such as node classification and clustering. This could be done either by fine-tuning CLEAR embeddings or retraining CLEAR framework with new downstream objectives. Because CLEAR generalizes well across multiple benchmark datasets, we believe it could also be used to contextualize LLM embeddings for other disease categories, such as cardiovascular and digestive system diseases.

To improve memory and computational efficiency, in the future we plan to explore less memory intensive methods like State Space Model (SSM) as an alternative to CLEAR’s attention-based modules. To enhance CLEAR’s interpretability, we will work on incorporating dedicated explainer module to clarify the reasoning behind every prediction. Finally, we aim to expand the ADRD KG by integrating additional data modalities, such as genomics and clinical data, to further enrich the context provided to CLEAR.

In summary, we believe our work demonstrates how to effectively leverage both the rich features from general-purpose biomedical LLMs and the structural information from biomedical KGs to enhance downstream computational tasks like novel drug-disease association prediction. As demonstrated in our application to complex diseases like ADRD, this framework may provide a valuable filtering tool for researchers seeking to discover effective treatments for diseases that currently lack or have limited therapeutic options.

## Methods

### CLEAR: A KG framework for contextualizing general-purpose LLM Embeddings

Pretrained LLMs are effective for generating high-dimensional feature representations of biomedical entities. However, their direct use in predictive modeling presents two key challenges. First, embeddings of different entities (e.g., drugs and diseases) are often generated by different LLMs, resulting in features that exist in separate, unaligned high-dimensional spaces, preventing direct comparison. Second, these embeddings are inherently general-purpose, as they are trained on vast corpuses of biomedical data, and may therefore lack the necessary context for a specific biomedical task. To overcome these limitations, we developed CLEAR (**C**ontextualizing **L**LM **E**mbeddings via **A**ttention-based g**R**aph learning), a framework that uses biomedical KG to contextualize general-purpose LLM embeddings.

We applied CLEAR framework to contextualize LLM embeddings specifically for Alzheimer Disease and Related Dementias (ADRD). The resulting contextualized embeddings were then used to identify and prioritize potential DR candidates for ADRD.

### KG construction and feature initialization

To represent multi-modal biological and pharmacological information about ADRD, we constructed a heterogeneous attributed KG, termed the Alzheimer Disease and Related Dementia KG (ADRD KG) (**Fig. 1a)**. Alzheimer disease and related dementias (ADRD) is an NIH/NIA research term encompassing Alzheimer disease, Lewy body dementias (including those associated with Parkinson disease), frontotemporal degeneration, vascular contributions to cognitive impairment and dementia, and mixed etiologies, reflecting converging biological and clinical pathways among neurodegenerative dementias but not constituting a formal DSM-5 or ICD-11 diagnostic category *per se*^105^. Because these neurodegenerative diseases are rooted within the CNS, drug candidates for AD and ADRD exhibit higher clinical failure rates and longer regulatory review times than non-CNS drugs^106^.

ADRD KG consists of three node types (drugs, diseases, and proteins) and six edge types representing relationships between them. Bipartite edges were sourced from public databases: drug-protein associations were obtained from DrugBank (v5.1.13)^107^, while drug–disease and disease–protein associations were from the Comparative Toxicogenomic Database (CTD 2025 release)^108^. Drug similarity edges and protein similarity edges were obtained from DrugSimDB^109^ and the STRING^110^ database, respectively. Disease similarity edges were computed based on Medical Subject Headings (MeSH) and Unified Medical Language System (UMLS)^111^ using the pyMeSHSim Python package ^112^. Node-specific attributes were also curated from these sources: drug SMILES strings from DrugBank, disease descriptions from MeSH, and protein sequences from UniProt. To make ADRD KG more disease-context specific, we only included U.S. Food and Drug Administration (FDA)-approved drugs, neuro-degenerative diseases classified under MeSH code C10 (Nervous System Diseases), and their respective associated therapeutic proteins as drug, disease, and protein nodes in ADRD KG, respectively.

Each node in the ADRD KG was initialized with a rich feature vector derived from a pretrained LLM specific to its data modality. For drug nodes, 768-dimensional embeddings were generated from their SMILES strings using MoLFormer. For disease nodes, the “Scope Note” or disease descriptions from the MeSH database were fed into BioBERT to produce 768-dimensional embeddings. Finally, for protein nodes, 1280-dimensional embeddings were generated from their amino acid sequences (obtained from UniProt) using ESM-2. This process resulted in a unique, high-dimensional feature representation for every node in the ADRD KG prior to the graph learning stage. To ensure data consistency for downstream analysis, all entity identifiers were standardized to DrugBank IDs (for drugs), MeSH IDs (for diseases), and UniProt IDs (for proteins).

### Mathematical formulation of ADRD KG and learning objective

CLEAR framework is built upon the ADRD KG, which integrates drug-related, disease-related and therapeutic protein related information. We formally represent this graph as *G*_*R*−*D*−*T*_ (*V*, *E*), where *V* constitutes the set of nodes and *E* represents the set of edges/relationships. The nodes in ADRD KG belong to one of three biomedical entries: drugs (*R*), diseases (*D*), and therapeutic proteins (*T*). The edges (*E*), which correspond to curated associations, are composed of three similarity networks (*G*_*R*_(*V*, *E*), *G*_*D*_(*V*, *E*), *G*_*T*_(*V*, *E*)) and three bipartite networks (*G*_*R*−*D*_(*V*, *E*), *G*_*R*−*T*_(*V*, *E*), *G*_*D*−*T*_(*V*, *E*)). Each node is initialized with a general-purpose feature vector obtained from a pretrained LLM, denoted as *x*_*R*_, *x*_*D*_ and *x*_*T*_ for drug, disease and protein nodes, respectively.

The primary objective of CLEAR is to learn a transformation that projects general-purpose LLM embedding into a single, unified embedding space. We achieve this by formulating the learning process for CLEAR as a supervised link prediction task. By training a model to predict the existence of edges within the ADRD KG, we train CLEAR to learn node representations (i.e., the CLEAR embeddings) such that the embeddings of pharmacologically and biologically related entities become more similar compared to similarities of their general-purpose LLM embeddings. These refined embeddings can then be used for downstream applications, such as identifying novel drug-disease associations for DR.

### Contextualizing embeddings via graph representation learning

The central module of CLEAR is a multi-relational GNN designed to learn from the complex, heterogeneous structure of the ADRD KG (**Fig. 5**). The goal of this module is to transform the initial LLM features into contextualized CLEAR embeddings. The following subsections detail the core architectural components of this GNN, which collectively produce a single, comprehensive representation for each node.

**Fig. 5:**
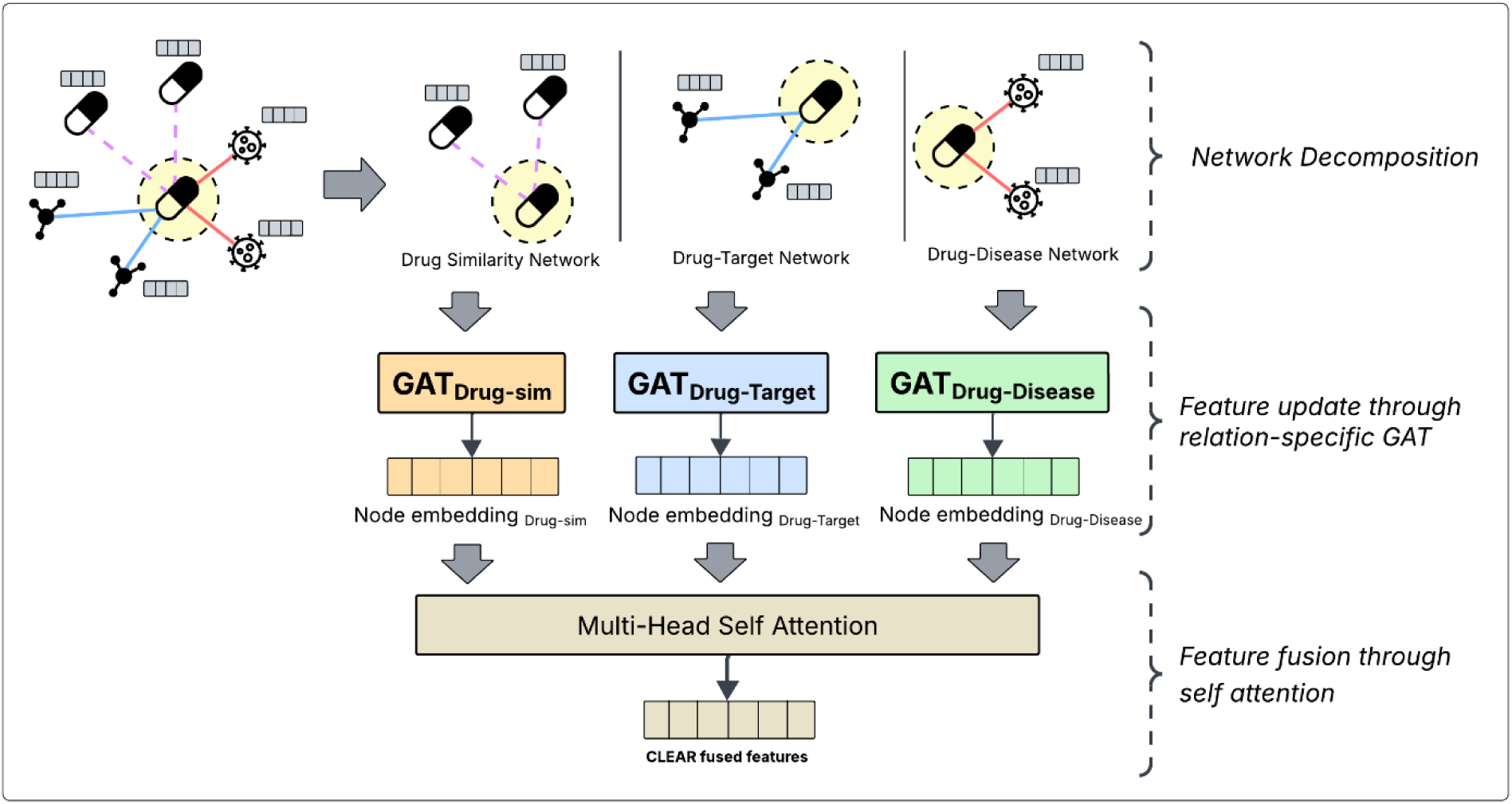
The CLEAR node embedding generation pipeline. The process involves three key stages. **a.** Network decomposition: The heterogeneous ADRD KG is first decomposed into its constituent homogeneous and bipartite subgraphs based on relation type. **b.** Relation-specific embedding: A dedicated Graph Attention Network (GAT) is then applied to each subgraph to generate relation-specific embeddings, preserving the unique semantics of each relation type. **c.** Embedding aggregation: Finally, these multiple, relation-specific embeddings are aggregated into a single, unified representation for each node using Multi-Head Self-Attention (MHSA).

### Initial Feature Transformation

The initial node features for drugs, diseases, and proteins were derived from different pretrained LLMs and thus had varying dimensionalities. To address this, we applied a linear transformation, which begins with a node-specific transformation that transforms the initial features of all node types into the same size. This was followed by a single transformation with a shared weight across all node types to project them into a unified embedding space (**Fig. 6**). This combined process can be represented by the following equation:

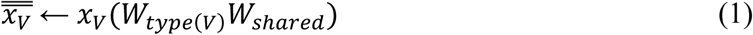

**Fig. 6.**
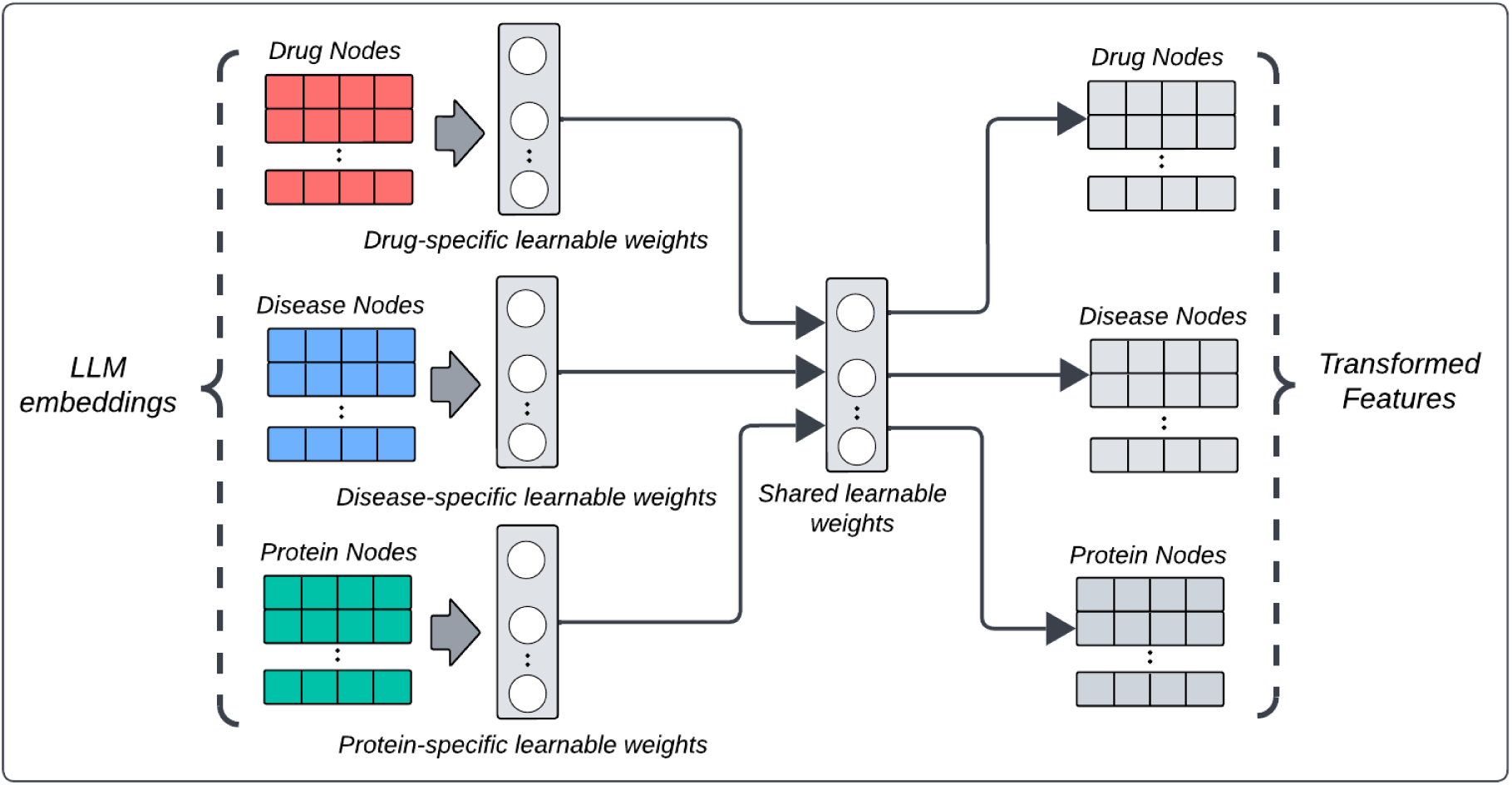
Initial Feature Transformation. To resolve dimensional inconsistencies, initial node features are projected into a unified embedding space. This is achieved in two steps: node type-specific layers first map features to a common dimension, followed by a final shared transformation layer.

In Equation 1, *x*_*V*_ and 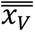 are the initial and transformed feature vector for a node of type *V* ∈ {*R*, *D*, *T*}, respectively. The term *W*_*type*(*V*)_ represents the node-type-specific weight matrix (W_R_, W_D_ or W_T_), and *W*_*shared*_ is the shared weight matrix applied across all node types.

### Generating relation-specific embeddings with GATs

After the initial node features are transformed into a unified space, the next step is to generate relation-specific node embeddings (**Supplementary Algorithm 2**). The core of our approach is to apply a separate Graph Attention Network (GAT) for each of the six subgraphs (three similarities and three bipartite networks). Applying separate GAT to individual networks helps preserve network-specific signals, effectively capturing the distinct structural characteristics inherent to each type of relationship. For instance, the signals that define drug-drug similarity are fundamentally different from those defining a drug-protein interaction. A single, shared GAT could cause these distinct signals to compete and become diluted, potentially losing important domain-specific information. We applied separate multi-head GATs initialized with same learnable parameters on each of the subgraphs.

For the three similarity subgraphs (*G*_*V*_, *where V* = {*R*, *D*, *T*}), three two-layer GATs were applied to generate relation-specific features as follows, where *f* is the *ReLU* activation function:

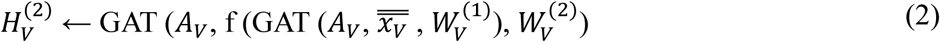

For the three bipartite subgraphs (*G*_*U*−*V*_, *where* (*U*, *V*) ∈ {(*R*, *D*), (*R* − *T*), (*D*, *T*)}, we first concatenated the feature vectors of the interacting node types. A two-layer GAT was then applied to these concatenated features:

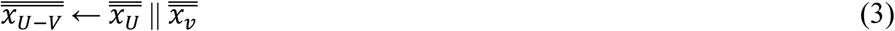

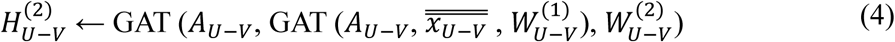

This process results in each node having multiple context-specific embeddings. For example, a drug node will have embeddings computed from the drug-drug similarity, drug-disease bipartite, and drug-protein bipartite network views. These embeddings are then ready for the final aggregation step.

### Fusing relation-specific embeddings with Multi-Head Self-Attention

Following the update of LLM embedding through multiple relation-specific GATs, we performed a fusion step to combine them into a single, comprehensive feature vector, which we named *CLEAR embedding*s. For example, the three separate embeddings for a drug node (from the drug-drug, drug-disease, and drug-protein views) are fused into one. This is achieved using Multi-Head Self-Attention (MHSA) mechanism, as detailed in **Supplementary Algorithm 3.**

The fusion process began by stacking the three relation-specific features for a given node into a single matrix, *X*. This matrix was then linearly projected into three separate matrices: Query (*Q*), Key (*K*) and Value (*V*) across multiple attention heads, using learnable weight matrices *W*_Q_, *W*_K_, *W*_V_, respectively:

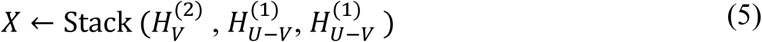

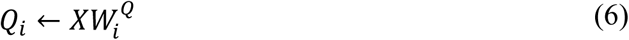

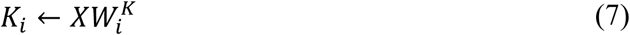

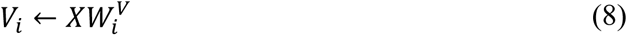

where *i* denotes the index of attention head

At each head the attention weights were computed by applying a SoftMax function (σ) to the scaled dot-product of the *Q* and *K* matrices. The output was a weighted sum of the *V* matrix vectors, based on these attention weights.:

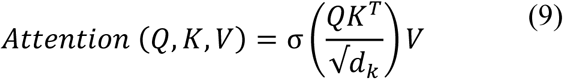

where *d*_*k*_ is the dimension of the *K* vectors.

The *Multi-Head* aspect of the mechanism performs this self-attention operation multiple times in parallel, each with different, independently learned weight matrices. This allows the model to jointly attend to information from different representation subspaces. The outputs of each head are then concatenated and passed through a final linear layer to produce the aggregated embedding, *Z*_*CLEAR*_:

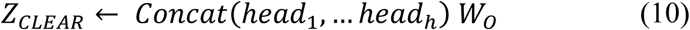

where *head*_*i*_ = *Attention* (*Q*_*i*_, *K*_*i*_, *V*_*i*_) and *W*_*o*_ is a final weight matrix to project the concatenated output back to embedding dimension.

This approach yields CLEAR embeddings 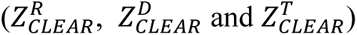 for drug, disease and protein, respectively that encapsulate information from multiple relational contexts and can be used for downstream tasks.

### Link prediction and model training

Once the CLEAR embeddings 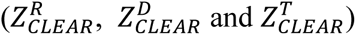 are computed, they can be used for the downstream task of link prediction. The prediction head is a MLP that functions as a binary classifier. As detailed in **Supplementary Algorithm 4**, the embeddings for any given pair of nodes (*U*, *V*) are first combined by concatenation to create a single vector representing the potential interaction:

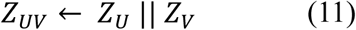

This combined vector was then fed into an MLP, which acted as a binary classifier to predict the existence of a link. The MLP outputs a final prediction score (*ŷ*_*UV*_), which represents the probability of an association between the two nodes:

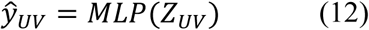

To address the imbalance between the sparse bipartite connections and the more frequent similarity links in the ADRD graph, we utilized a weighted binary cross-entropy loss function to update model parameters. This prevents the model from becoming biased toward the abundant similarity links during training. The overall loss, *L*, is defined as:

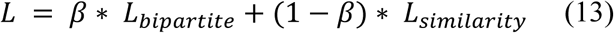

Here, *β* is a predefined hyperparameter that controls the trade-off, giving higher importance to the bipartite links. The term *L*_*bipartite*_ is the average binary cross-entropy loss over the three bipartite link types (i.e., drug-disease, drug-protein, and disease-protein). Similarly, *L*_*similarity*_ is the average loss over the three similarity link types (i.e., drug-drug, disease-disease, and protein-protein).

### Topology-aware negative sampling

To train CLEAR for link prediction, we formalize the task by labeling node pairs. Any (source node, destination node) pair connected by an edge in the original ADRD KG is considered a positive instance and labeled as ‘**1**’. To generate negative instances labeled as ‘**0**’, we employed a topology-aware negative sampling strategy with three key principles, as detailed in **Supplementary Algorithm 1**. First, to avoid sampling false negatives, we generate negative pairs for a source node by exclusively sampling destination nodes that lie outside of its 3-hop neighborhood. Second, to create challenging negative examples, we ensure the node degree distribution of the sampled negative pairs is like that of the positive pairs. This forces the model to learn complex structural patterns beyond immediate topological information. Third, these sampling rules are applied independently to each similarity and bipartite subgraphs (*G*_*R*_(*V*, *E*), *G*_*D*_(*V*, *E*), *G*_*T*_(*V*, *E*), *G*_*R*−*D*_(*V*, *E*), *G*_*R*−*T*_(*V*, *E*), *G*_*D*−*T*_(*V*, *E*)). This approach allows all associations to have equal representation during training and testing, preventing scenarios where certain types of associations disproportionately dominate any single dataset partition.

### KG completion and candidate drug prioritization

Once CLEAR framework is trained, we reconstruct the ADRD KG to generate comprehensive map of potential interactions. This reconstruction applies only to the bipartite links such as drug-disease associations, drug-protein interactions, and disease-protein associations. For similarity links, such as drug-drug, disease-disease, and protein-protein connections, we retain the existing known connections and do not predict new links. To predict new bipartite links, we generate all pairwise CLEAR embedding combinations of all drugs, disease and protein nodes and fed into pretrained link prediction model (MLP) to generate link probability score. For a given node pair, if link probability score is greater than 0.5, a new edge is added between them in ADRD KG. Once all possible edges are processed, we end up with updated ADRD KG.

Following the reconstruction of ADRD KG, we prioritize potential *Drug candidates* for a given query *Disease* using multi-criteria ranking approach as follows:

a. Confidence of predicted drug-disease link: Drugs of predicted drug-disease links with high link probability are ranked higher as these indicates a stronger association.
b. Therapeutic target overlap: We assess the degree of overlap between the known therapeutic targets of a drug and the targets associated with the query disease^113,114^. Drugs are ranked based on the extent of this shared target module, rewarding candidates that are mechanistically relevant to the disease’s biology.

Drugs that are ranked high across both criteria are considered as *Drug candidates* for user queried *Disease*.

## References

1. Singh, N., Vayer, P., Tanwar, S., Poyet, J.-L., Tsaioun, K., & Villoutreix, B. O. (2023). Drug discovery and development: introduction to the general public and patient groups. Frontiers in Drug Discovery, 3. 10.3389/fddsv.2023.1201419

2. DiMasi, J. A. (2020). Research and Development Costs of New Drugs. JAMA, 324(5), 517. 10.1001/jama.2020.8648

3. Scott, T. J., O’Connor, A. C., Link, A. N., & Beaulieu, T. J. (2014). Economic analysis of opportunities to accelerate Alzheimer’s disease research and development. Annals of the New York Academy of Sciences, 1313(1), 17–34. 10.1111/nyas.12417

4. Krishnamurthy, N., Grimshaw, A. A., Axson, S. A., Choe, S. H., & Miller, J. E. (2022). Drug repurposing: a systematic review on root causes, barriers and facilitators. BMC Health Services Research, 22(1). 10.1186/s12913-022-08272-z

5. Ashburn, T. T., & Thor, K. B. (2004). Drug repositioning: Identifying and developing new uses for existing drugs. Nature Reviews Drug Discovery, 3(8), 673–683. 10.1038/nrd1468

6. Wallach, Izhar, et al. “AI Is a Viable Alternative to High Throughput Screening: A 318-Target Study.” Scientific Reports, vol. 14, no. 1, 2 Apr. 2024, p. 7526, www.nature.com/articles/s41598-024-54655-z, 10.1038/s41598-024-54655-z.

7. Surabhi, S., & Singh, B. (2018). Computer aided drug design: an overview. J. Drug Deliv. Ther, 8(5), 504–509

8. Oliveira, T. A. D., Silva, M. P. D., Maia, E. H. B., Silva, A. M. D., & Taranto, A. G. (2023). Virtual screening algorithms in drug discovery: a review focused on machine and deep learning methods. Drugs and Drug Candidates, 2(2), 311–334. 10.3390/ddc2020017

9. Vitorino, R. (2024). Transforming Clinical Research: The Power of High-Throughput Omics Integration. Proteomes, 12(3), 25–25. 10.3390/proteomes12030025

10. Cui, M., Cheng, C., & Zhang, L. (2022). High-throughput proteomics: A methodological mini-review. Laboratory Investigation; a Journal of Technical Methods and Pathology, 102(11), 1170–1181. 10.1038/s41374-022-00830-7

11. Jarada, Tamer N., et al. “A Review of Computational Drug Repositioning: Strategies, Approaches, Opportunities, Challenges, and Directions.” Journal of Cheminformatics, vol. 12, no. 1, 22 July 2020, 10.1186/s13321-020-00450-7.

12. Wang, Yongcui, et al. “Drug Repositioning by Kernel-Based Integration of Molecular Structure, Molecular Activity, and Phenotype Data.” PLoS ONE, vol. 8, no. 11, 11 Nov. 2013, p. e78518, 10.1371/journal.pone.0078518. Accessed 17 Mar. 2023.

13. Yang, Lun, and Pankaj Agarwal. “Systematic Drug Repositioning Based on Clinical Side-Effects.” PLoS ONE, vol. 6, no. 12, 21 Dec. 2011, p. e28025, 10.1371/journal.pone.0028025. Accessed 20 Oct. 2020.

14. Yang, M., Luo, H., Li, Y., & Wang, J. (2019). Drug repositioning based on bounded nuclear norm regularization. Bioinformatics, 35(14), i455–i463. 10.1093/bioinformatics/btz331

15. Zhang, W., Yue, X., Lin, W., Wu, W., Liu, R., Huang, F., & Liu, F. (2018). Predicting drug-disease associations by using similarity constrained matrix factorization. BMC Bioinformatics, 19(1). 10.1186/s12859-018-2220-4

16. Zhang, W., Xu, H., Li, X., Gao, Q., & Wang, L. (2020). DRIMC: an improved drug repositioning approach using Bayesian inductive matrix completion. Bioinformatics, 36(9), 2839–2847. 10.1093/bioinformatics/btaa062

17. Luo, H., Wang, J., Li, M., Luo, J., Peng, X., Wu, F.-X., & Pan, Y. (2016). Drug repositioning based on comprehensive similarity measures and Bi-Random walk algorithm. Bioinformatics, 32(17), 2664–2671. 10.1093/bioinformatics/btw228

18. Chen, H., Cheng, F., & Li, J. (2020). iDrug: Integration of drug repositioning and drug-target prediction via cross-network embedding. PLOS Computational Biology, 16(7), e1008040. 10.1371/journal.pcbi.1008040

19. Bang, D., Lim, S., Lee, S., & Kim, S. (2023). Biomedical knowledge graph learning for drug repurposing by extending guilt-by-association to multiple layers. Nature Communications, 14(1), 3570. 10.1038/s41467-023-39301-y

20. Lijun Cai, Changcheng Lu, Junlin Xu, Yajie Meng, Peng Wang, Xiangzheng Fu, Xiangxiang Zeng, Yansen Su, Drug repositioning based on the heterogeneous information fusion graph convolutional network, Briefings in Bioinformatics, Volume 22, Issue 6, November 2021, bbab319, 10.1093/bib/bbab319

21. Yajie Meng, Yi Wang, Junlin Xu, Changcheng Lu, Xianfang Tang, Tao Peng, Bengong Zhang, Geng Tian, Jialiang Yang, Drug repositioning based on weighted local information augmented graph neural network, Briefings in Bioinformatics, Volume 25, Issue 1, January 2024, bbad431, 10.1093/bib/bbad431

22. Yan, Y., Yang, Y., Tong, Z., Wang, Y., Yang, F., Pan, Z., Liu, C., Bai, M., Xie, Y., Li, Y., Shu, K., & Li, Y. (2025). Adaptive multi-view learning method for enhanced drug repurposing using chemical-induced transcriptional profiles, knowledge graphs, and large language models. Journal of Pharmaceutical Analysis, 15(6), 101275. 10.1016/j.jpha.2025.101275

23. Wang, B., Xie, Q., Pei, J., Chen, Z., Tiwari, P., Zhao, L., & Fu, J. (2023). Pre-trained Language Models in Biomedical Domain: A Systematic Survey. ACM Computing Surveys, 56(3). 10.1145/3611651

24. Jia, Z.-C., Yang, X., Wu, Y.-K., Li, M., Das, D., Chen, M.-X., & Wu, J. (2024). The Art of Finding the Right Drug Target: Emerging Methods and Strategies. Pharmacological Reviews, 76(5), 896–914. 10.1124/pharmrev.123.001028

25. Lahiri, A., Shukla, S., Stear, B., Ahooyi, T. M., Beigel, K., Margolskee, E., & Taylor, D. (2025). Benchmarking transformer embedding models for biomedical terminology standardization. Machine Learning with Applications, 21, 100683. 10.1016/j.mlwa.2025.100683

26. Jarada, T. N., Rokne, J. G., & Alhajj, R. (2020). A review of computational drug repositioning: strategies, approaches, opportunities, challenges, and directions. Journal of Cheminformatics, 12(1). 10.1186/s13321-020-00450-7

27. Montine, Thomas J., et al. “Recommendations of the Alzheimer’s Disease–Related Dementias Conference.” Neurology, vol. 83, no. 9, 26 Aug. 2014, pp. 851–860, www.ncbi.nlm.nih.gov/pmc/articles/PMC4155046/, 10.1212/WNL.0000000000000733. Accessed 1 Mar. 2021.

28. Ross, J., Belgodere, B., Vijil Chenthamarakshan, Inkit Padhi, Youssef Mroueh, & Das, P. (2022). Molformer: Large Scale Chemical Language Representations Capture Molecular Structure and Properties. Research Square (Research Square). 10.21203/rs.3.rs-1570270/v1

29. Lee, J., Yoon, W., Kim, S., Kim, D., Kim, S., So, C. H., & Kang, J. (2019). BioBERT: a pre-trained biomedical language representation model for biomedical text mining. Bioinformatics, 36(4). 10.1093/bioinformatics/btz682

30. Lin, Z., Akin, H., Rao, R., Hie, B., Zhu, Z., Lu, W., Smetanin, N., Verkuil, R., Kabeli, O., Shmueli, Y., dos Santos Costa, A., Fazel-Zarandi, M., Sercu, T., Candido, S., & Rives, A. (2023). Evolutionary-scale prediction of atomic-level protein structure with a language model. Science, 379(6637), 1123–1130. 10.1126/science.ade2574

31. Gottlieb, A., Stein, G. Y., Ruppin, E., & Sharan, R. (2011). PREDICT: a method for inferring novel drug indications with application to personalized medicine. Molecular Systems Biology, 7, 496. 10.1038/msb.2011.26

32. Zhouxin Yu, Feng Huang, Xiaohan Zhao, Wenjie Xiao, Wen Zhang, Predicting drug–disease associations through layer attention graph convolutional network, Briefings in Bioinformatics, Volume 22, Issue 4, July 2021, bbaa243, 10.1093/bib/bbaa243

33. Yang, M., Wu, G., Zhao, Q., Li, Y., & Wang, J. (2020). Computational drug repositioning based on multi-similarities bilinear matrix factorization. Briefings in Bioinformatics, 22(4). 10.1093/bib/bbaa267

34. Liang, X., Zhang, P., Yan, L., Fu, Y., Peng, F., Qu, L., Shao, M., Chen, Y., & Chen, Z. (2016). LRSSL: predict and interpret drug–disease associations based on data integration using sparse subspace learning. Bioinformatics, 33(8), 1187–1196. 10.1093/bioinformatics/btw770

35. Davis, A., Wiegers, T., Sciaky, D., Barkalow, F., Strong, M., Wyatt, B., Wiegers, J., McMorran, R., Sakib Abrar, & Mattingly, C. (2024). Comparative toxicogenomics database’s 20th anniversary: update 2025. Nucleic Acids Research, 53(D1), D1328–D1334. 10.1093/nar/gkae883

36. Li, Y., Yang, Y., Tong, Z., Wang, Y., Mi, Q., Bai, M., Liang, G., Li, B., & Shu, K. (2024). A comparative benchmarking and evaluation framework for heterogeneous network-based drug repositioning methods. Briefings in Bioinformatics, 25(3). 10.1093/bib/bbae172

37. Yang, M., Yang, B., Duan, G., & Wang, J. (2023). ITRPCA: a new model for computational drug repositioning based on improved tensor robust principal component analysis. Frontiers in Genetics, 14. 10.3389/fgene.2023.1271311

38. Yan, Y., Yang, M., Zhao, H., Duan, G., Peng, X., & Wang, J. (2022). Drug repositioning based on multi-view learning with matrix completion. Briefings in Bioinformatics, 23(3). 10.1093/bib/bbac054

39. Yang, M., Luo, H., Li, Y., Wu, F.-X., & Wang, J. (2019). Overlap matrix completion for predicting drug-associated indications. PLOS Computational Biology, 15(12), e1007541. 10.1371/journal.pcbi.1007541

40. Wang, Y., Xiang, J., Liu, C., Tang, M., Hou, R., Bao, M., Tian, G., He, J., & He, B. (2022). Drug repositioning for SARS-CoV-2 by Gaussian kernel similarity bilinear matrix factorization. Frontiers in Microbiology, 13. 10.3389/fmicb.2022.1062281

41. Zhao, B.-W., Hu, L., You, Z.-H., Wang, L., & Su, X. (2021). HINGRL: predicting drug–disease associations with graph representation learning on heterogeneous information networks. Briefings in Bioinformatics, 23(1). 10.1093/bib/bbab515

42. Luo, H., Li, M., Wang, S., Liu, Q., Li, Y., & Wang, J. (2018). Computational drug repositioning using low-rank matrix approximation and randomized algorithms. Bioinformatics, 34(11), 1904–1912. 10.1093/bioinformatics/bty013

43. Xie, G., Xu, H., Li, J., Gu, G., Sun, Y., Lin, Z., Zhu, Y., Wang, W., Wang, Y., & Shao, J. (2022). DRPADC: A novel drug repositioning algorithm predicting adaptive drugs for COVID-19. Computers & Chemical Engineering, 166, 107947–107947. 10.1016/j.compchemeng.2022.107947

44. Yang, M., Huang, L., Xu, Y., Lu, C., & Wang, J. (2020). Heterogeneous graph inference with matrix completion for computational drug repositioning. Bioinformatics (Oxford. Print), 36(22-23), 5456–5464. 10.1093/bioinformatics/btaa1024

45. Mongia, A., Chouzenoux, E., & Majumdar, A. (2022). Computational Prediction of Drug-Disease Association Based on Graph-Regularized One Bit Matrix Completion. IEEE/ACM Transactions on Computational Biology and Bioinformatics, 19(6), 3332–3339. 10.1109/tcbb.2022.3189879

46. Wang, X., & Yan, R. (2020). DDAPRED: a computational method for predicting drug repositioning using regularized logistic matrix factorization. Journal of Molecular Modeling, 26(3). 10.1007/s00894-020-4315-x

47. Watt, N. T., Whitehouse, I. J., & Hooper, N. M. (2011). The Role of Zinc in Alzheimer’s Disease. International Journal of Alzheimer’s Disease, 2011, 1–10. 10.4061/2011/971021

48. Bush, A., Pettingell, W., Multhaup, G., d Paradis, M., Vonsattel, J., Gusella, J., Beyreuther, K., Masters, C., & Tanzi, R. (1994). Rapid induction of Alzheimer A beta amyloid formation by zinc. Science, 265(5177), 1464–1467. 10.1126/science.8073293

49. Craddock, T. J. A., Tuszynski, J. A., Chopra, D., Casey, N., Goldstein, L. E., Hameroff, S. R., & Tanzi, R. E. (2012). The Zinc Dyshomeostasis Hypothesis of Alzheimer’s Disease. PLoS ONE, 7(3), e33552. 10.1371/journal.pone.0033552

50. Schrag, M., Mueller, C., Oyoyo, U., Smith, M. A., & Kirsch, W. M. (2011). Iron, zinc and copper in the Alzheimer’s disease brain: A quantitative meta-analysis. Some insight on the influence of citation bias on scientific opinion. Progress in Neurobiology, 94(3), 296–306. 10.1016/j.pneurobio.2011.05.001

51. Sabalic, A., Mei, V., Giuliana Solinas, & Madeddu, R. (2024). The Role of Copper in Alzheimer’s Disease Etiopathogenesis: An Updated Systematic Review. Toxics, 12(10), 755–755. 10.3390/toxics12100755

52. Taylor, C. P., Traynelis, S. F., Siffert, J., Pope, L. E., & Matsumoto, R. R. (2016). Pharmacology of dextromethorphan: Relevance to dextromethorphan/quinidine (Nuedexta®) clinical use. Pharmacology & Therapeutics, 164, 170–182. 10.1016/j.pharmthera.2016.04.010

53. Chen, C.-Y., Chung, C.-H., Chien, W.-C., & Chen, H.-C. (2022). The Association Between Dextromethorphan Use and the Risk of Dementia. American Journal of Alzheimer’s Disease & Other Dementias®, 37, 153331752211249. 10.1177/15333175221124952

54. Lin, L., Huang, Q.-X., Yang, S.-S., Chu, J., Wang, J.-Z., & Tian, Q. (2013). Melatonin in Alzheimer’s Disease. International Journal of Molecular Sciences, 14(7), 14575–14593. 10.3390/ijms140714575

55. Olivieri, G., Hess, C., Savaskan, E., Ly, C., Meier, F., Baysang, G., Brockhaus, M., & Müller-Spahn, F. (2001). Melatonin protects SHSY5Y neuroblastoma cells from cobalt-induced oxidative stress, neurotoxicity and increased β-amyloid secretion. Journal of Pineal Research, 31(4), 320–325. 10.1034/j.1600-079x.2001.310406.x

56. Ebrahimi, R., Faramarzi, A., Salarvandian, S., Zarei, R., Heidari, M., Salehian, F., & Esmaeilpour, K. (2025). Melatonin Supplementation in Alzheimer’s disease: The Potential Role in Neurogenesis. Molecular Neurobiology. 10.1007/s12035-025-05095-x

57. Coelho Filho, J. M. J., & Birks, J. (2001). Physostigmine for dementia due to Alzheimer’s disease. Cochrane Database of Systematic Reviews. 10.1002/14651858.cd001499

58. Ramesh, S, and S Perera. “Depletion of Dopamine in Parkinson’s Disease and Relevant Therapeutic Options: A Review of the Literature.” AIMS Neuroscience, vol. 10, no. 3, 1 Jan. 2023, pp. 200–231, pmc.ncbi.nlm.nih.gov/articles/PMC10567584/, 10.3934/neuroscience.2023017.

59. Latif, Saad, et al. “Dopamine in Parkinson’s Disease.” Clinica Chimica Acta, vol. 522, no. 0009-8981, 2021, pp. 114–126, www.sciencedirect.com/science/article/pii/S000989812100276X?via%3Dihub, 10.1016/j.cca.2021.08.009.

60. Davies, Katherine M., et al. “Copper Dyshomoeostasis in Parkinson’s Disease: Implications for Pathogenesis and Indications for Novel Therapeutics.” Clinical Science, vol. 130, no. 8, 8 Mar. 2016, pp. 565–574, portlandpress.com/clinsci/article/130/8/565/71505/Copper-dyshomoeostasis-in-Parkinson-s-disease, 10.1042/cs20150153.

61. Montes, Sergio, et al. “Copper and Copper Proteins in Parkinson’s Disease.” Oxidative Medicine and Cellular Longevity, vol. 2014, 2014, pp. 1–15, www.ncbi.nlm.nih.gov/pmc/articles/PMC3941957/, 10.1155/2014/147251.

62. Wang, Hai-Li, et al. “Potential Use of Glutathione as a Treatment for Parkinson’s Disease.” Experimental and Therapeutic Medicine, vol. 21, no. 2, 4 Dec. 2020, 10.3892/etm.2020.9557.

63. Hauser, Robert A., et al. “Randomized, Double-Blind, Pilot Evaluation of Intravenous Glutathione in Parkinson’s Disease.” Movement Disorders, vol. 24, no. 7, 19 Feb. 2009, pp. 979–983, 10.1002/mds.22401.

64. Smeyne, Michelle, and Richard Jay Smeyne. “Glutathione Metabolism and Parkinson’s Disease.” Free Radical Biology and Medicine, vol. 62, Sept. 2013, pp. 13–25, www.ncbi.nlm.nih.gov/pmc/articles/PMC3736736/, 10.1016/j.freeradbiomed.2013.05.001.

65. Forsleff, Louise, et al. “Evidence of Functional Zinc Deficiency in Parkinson’s Disease.” The Journal of Alternative and Complementary Medicine, vol. 5, no. 1, Feb. 1999, pp. 57–64, 10.1089/acm.1999.5.57.

66. Saini, Nidhi, and Walter Schaffner. “Zinc Supplement Greatly Improves the Condition of Parkin Mutant Drosophila.” Biological Chemistry, vol. 391, no. 5, 1 May 2010, pp. 513–518, 10.1515/bc.2010.052.

67. Brewer, George J., et al. “Subclinical Zinc Deficiency in Alzheimer’s Disease and Parkinson’s Disease.” American Journal of Alzheimer’s Disease & Other Dementias, vol. 25, no. 7, 14 Sept. 2010, pp. 572–575, 10.1177/1533317510382283.

68. Wu, Leslie, and Archana Jhawar. “Loxapine for the Treatment of Psychosis in Lewy Body Dementia: A Case Report.” Mental Health Clinician, vol. 15, no. 5, Oct. 2025, pp. 252–254, 10.9740/mhc.2025.10.252.

69. Lin, Chih-Li, et al. “Amitriptyline Improves Cognitive and Neuronal Function in a Rat Model That Mimics Dementia with Lewy Bodies.” Behavioural Brain Research, vol. 435, 28 Oct. 2022, p. 114035, www.sciencedirect.com/science/article/pii/S0166432822003035, 10.1016/j.bbr.2022.114035.

70. Ali, Noor, et al. “The Role of Estrogen Therapy as a Protective Factor for Alzheimer’s Disease and Dementia in Postmenopausal Women: A Comprehensive Review of the Literature.” Cureus, vol. 15, no. 8, 6 Aug. 2023, 10.7759/cureus.43053.

71. Whitfield, David R., et al. “Depression and Synaptic Zinc Regulation in Alzheimer Disease, Dementia with Lewy Bodies, and Parkinson Disease Dementia.” The American Journal of Geriatric Psychiatry, vol. 23, no. 2, Feb. 2015, pp. 141–148, 10.1016/j.jagp.2014.05.001. Accessed 15 Dec. 2021.

72. Varidaki, Artemis, et al. “Repositioning Microtubule Stabilizing Drugs for Brain Disorders.” Frontiers in Cellular Neuroscience, vol. 12, 8 Aug. 2018, 10.3389/fncel.2018.00226.

73. Patterson, Joseph R., et al. “Beta2-Adrenoreceptor Agonist Clenbuterol Produces Transient Decreases in Alpha-Synuclein MRNA but No Long-Term Reduction in Protein.” Npj Parkinson’s Disease, vol. 8, no. 1, 24 May 2022, 10.1038/s41531-022-00322-x.

74. Church, John, et al. “Interactions of Dextromethorphan with TheN-Methyl-d-Aspartate Receptor-Channel Complex: Single Channel Recordings.” Brain Research, vol. 666, no. 2, Dec. 1994, pp. 189–194, 10.1016/0006-8993(94)90771-4. Accessed 25 Mar. 2021.

75. Siu, Anita, and Richard Drachtman. “Dextromethorphan: A Review of N-Methyl-d-Aspartate Receptor Antagonist in the Management of Pain.” CNS Drug Reviews, vol. 13, no. 1, Mar. 2007, pp. 96–106, 10.1111/j.1527-3458.2007.00006.x.

76. Olivares, David, et al. “N-Methyl D-Aspartate (NMDA) Receptor Antagonists and Memantine Treatment for Alzheimer’s Disease, Vascular Dementia and Parkinson’s Disease.” Current Alzheimer Research, vol. 9, no. 6, 2012, pp. 746–58, www.ncbi.nlm.nih.gov/pubmed/21875407, 10.2174/156720512801322564.

77. McShane, Rupert, et al. “Memantine for Dementia.” Cochrane Database of Systematic Reviews, vol. 3, no. 3, 20 Mar. 2019, www.cochrane.org/CD003154/DEMENTIA_memantine-treatment-dementia, 10.1002/14651858.cd003154.pub6.

78. Shin, E, et al. “Dextromethorphan Attenuates Trimethyltin-Induced Neurotoxicity via σ1 Receptor Activation in Rats.” Neurochemistry International, vol. 50, no. 6, 5 Feb. 2007, pp. 791–799, 10.1016/j.neuint.2007.01.008. Accessed 15 July 2025.

79. Nguyen, Linda, et al. “Dextromethorphan: An Update on Its Utility for Neurological and Neuropsychiatric Disorders.” Pharmacology & Therapeutics, vol. 159, Mar. 2016, pp. 1–22, 10.1016/j.pharmthera.2016.01.016.

80. Eskandari, Kiarash, et al. “Repurposing Sigma-1 Receptor-Targeting Drugs for Therapeutic Advances in Neurodegenerative Disorders.” Pharmaceuticals, vol. 18, no. 5, 9 May 2025, p. 700, scite.ai/reports/repurposing-sigma-1-receptor-targeting-drugs-for-0GPzWe5Z, 10.3390/ph18050700. Accessed 15 May 2025.

81. Sethi, Roopa, et al. “Serotonin Syndrome in a Sertraline-Treated Man Taking NyQuil Containing Dextromethorphan for Cold.” The Primary Care Companion for CNS Disorders, vol. 14, no. 6, 2012, p. PCC.12l01388, www.ncbi.nlm.nih.gov/pmc/articles/PMC3622531/, 10.4088/PCC.12l01388. Accessed 25 Jan. 2022.

82. Singh, Michelle Anjali, and Devon Johnson. “Serotonin Syndrome and Dextromethorphan Toxicity Caused by Drug-Drug Interaction between Fluoxetine and Bupropion-Dextromethorphan: A Case Report.” The Journal of Clinical Psychiatry, vol. 85, no. 2, 3 Apr. 2024, p. 54018, www.psychiatrist.com/jcp/serotonin-syndrome-dextromethorphan-toxicity-with-fluoxetine-bupropion-dextromethorphan-interaction/, 10.4088/JCP.23cr15139.

83. Lee, Daniel, et al. “Brexpiprazole for Agitation Associated with Dementia due to Alzheimer’s Disease.” Journal of the American Medical Directors Association, 22 July 2024, p. 105173, pubmed.ncbi.nlm.nih.gov/39053890/, 10.1016/j.jamda.2024.105173.

84. Aga, Vimal M. “Brexpiprazole for the Treatment of Agitation in Alzheimer’s Disease Dementia: Clinical Uncertainties and the Path Forward.” The American Journal of Geriatric Psychiatry, 19 Nov. 2024, www.sciencedirect.com/science/article/abs/pii/S1064748124005335, 10.1016/j.jagp.2024.11.003.

85. Hernandez SC, Bertolino M, Xiao Y, Pringle KE, Caruso FS, Kellar KJ. Dextromethorphan and its metabolite dextrorphan block alpha3beta4 neuronal nicotinic receptors. J Pharmacol Exp Ther. 2000 Jun;293(3):962–7. PMID: 10869398.

86. Wrightjr, M, et al. “Comparative Effects of Dextromethorphan and Dextrorphan on Nicotine Discrimination in Rats.” Pharmacology Biochemistry and Behavior, vol. 85, no. 3, Nov. 2006, pp. 507–513, 10.1016/j.pbb.2006.09.020. Accessed 13 Feb. 2023.

87. Iosifescu, Dan V., et al. “Efficacy and Safety of AXS-05 (Dextromethorphan-Bupropion) in Patients with Major Depressive Disorder: A Phase 3 Randomized Clinical Trial (GEMINI).” The Journal of Clinical Psychiatry, vol. 83, no. 4, 30 May 2022, p. 41226, www.psychiatrist.com/jcp/depression/efficacy-safety-of-axs-05-dextromethorphan-bupropion-mdd/, 10.4088/JCP.21m14345.

88. Echeverria, V., Mendoza, C., & Iarkov, A. (2023). Nicotinic acetylcholine receptors and learning and memory deficits in Neuroinflammatory diseases. Frontiers in Neuroscience (Online), 17. 10.3389/fnins.2023.1179611

89. Geerts, H. (2011). α7 Nicotinic receptor modulators for cognitive deficits in schizophrenia and Alzheimer’s disease. Expert Opinion on Investigational Drugs, 21(1), 59–65. 10.1517/13543784.2012.633510

90. Vallés, A. S., Borroni, M. V., & Barrantes, F. J. (2014). Targeting Brain α7 Nicotinic Acetylcholine Receptors in Alzheimer’s Disease: Rationale and Current Status. CNS Drugs, 28(11), 975–987. 10.1007/s40263-014-0201-3

91. Izzo, N. J., Xu, J., Zeng, C., Kirk, M. J., Mozzoni, K., Silky, C., Rehak, C., Yurko, R., Look, G. C., Rishton, G. M., Safferstein, H., Cruchaga, C., Goate, A., Cahill, M. A., Arancio, O., Mach, R. H., Craven, R. J., Head, E., LeVine, H., & Spires-Jones, T. L. (2014b). Alzheimer’s Therapeutics Targeting Amyloid Beta 1–42 Oligomers II: Sigma-2/PGRMC1 Receptors Mediate Abeta 42 Oligomer Binding and Synaptotoxicity. PLOS ONE, 9(11), e111899–e111899. 10.1371/journal.pone.0111899

92. Riad, A., Zsofia Lengyel-Zhand, Zeng, C., Weng, C.-C., Virginia M.-Y. Lee, Trojanowski, J. Q., & Mach, R. H. (2020b). The Sigma-2 Receptor/TMEM97, PGRMC1, and LDL Receptor Complex Are Responsible for the Cellular Uptake of Aβ42 and Its Protein Aggregates. Molecular Neurobiology, 57(9), 3803–3813. 10.1007/s12035-020-01988-1

93. van Dyck, Christopher H., et al. “Lecanemab in Early Alzheimer’s Disease.” New England Journal of Medicine, vol. 388, no. 1, 29 Nov. 2022, pp. 9–21, www.nejm.org/doi/full/10.1056/NEJMoa2212948, 10.1056/nejmoa2212948.

94. Sims, John R., et al. “Donanemab in Early Symptomatic Alzheimer Disease: The TRAILBLAZER-ALZ 2 Randomized Clinical Trial.” JAMA, vol. 330, no. 6, 17 July 2023, jamanetwork.com/journals/jama/fullarticle/2807533, 10.1001/jama.2023.13239.

95. Budd Haeberlein, S., et al. “Two Randomized Phase 3 Studies of Aducanumab in Early Alzheimer’s Disease.” The Journal of Prevention of Alzheimer’s Disease, vol. 9, no. 2, 2022, pubmed.ncbi.nlm.nih.gov/35542991/, 10.14283/jpad.2022.30.

96. Simpson, D. S. A., & Oliver, P. L. (2020). ROS Generation in Microglia: Understanding Oxidative Stress and Inflammation in Neurodegenerative Disease. Antioxidants, 9(8), 743. 10.3390/antiox9080743

97. Wilkinson, B. L., & Landreth, G. E. (2006). The microglial NADPH oxidase complex as a source of oxidative stress in Alzheimer’s disease. Journal of Neuroinflammation, 3, 30. 10.1186/1742-2094-3-30

98. H. Ferreira-Vieira, T., M. Guimaraes, I., R. Silva, F., & M. Ribeiro, F. (2016). Alzheimer’s disease: Targeting the Cholinergic System. Current Neuropharmacology, 14(1), 101–115. 10.2174/1570159x13666150716165726

99. Maurer, S. V., & Williams, C. L. (2017). The Cholinergic System Modulates Memory and Hippocampal Plasticity via Its Interactions with Non-Neuronal Cells. Frontiers in Immunology, 8. 10.3389/fimmu.2017.01489

100. Chen, Chia-Yuan, et al. “The Association between Dextromethorphan Use and the Risk of Dementia.” American Journal of Alzheimer’s Disease & Other Dementias®, vol. 37, Jan. 2022, p. 153331752211249, 10.1177/15333175221124952.

101. Cummings, Jeffrey L., et al. “Effect of Dextromethorphan-Quinidine on Agitation in Patients with Alzheimer Disease Dementia.” JAMA, vol. 314, no. 12, 22 Sept. 2015, p. 1242, 10.1001/jama.2015.10214.

102. Gray, Shelly L, et al. “Cumulative Use of Strong Anticholinergics and Incident Dementia: A Prospective Cohort Study.” JAMA Internal Medicine, vol. 175, no. 3, 2015, pp. 401–7, 10.1001/jamainternmed.2014.7663.

103. Campbell, N. L., et al. “Use of Anticholinergics and the Risk of Cognitive Impairment in an African American Population.” Neurology, vol. 75, no. 2, 12 July 2010, pp. 152–159, 10.1212/wnl.0b013e3181e7f2ab.

104. Ruxton, Kimberley, et al. “Drugs with Anticholinergic Effects and Cognitive Impairment, Falls and All-Cause Mortality in Older Adults: A Systematic Review and Meta-Analysis.” British Journal of Clinical Pharmacology, vol. 80, no. 2, 20 May 2015, pp. 209–220, 10.1111/bcp.12617.

105. Huang, Z., Xiao, Z., Ao, C., Guan, L., & Yu, L. (2024). Computational approaches for predicting drug-disease associations: a comprehensive review. Frontiers of Computer Science, 19(5). 10.1007/s11704-024-40072-y

106. Gribkoff, V. K., & Kaczmarek, L. K. (2017). The need for new approaches in CNS drug discovery: Why drugs have failed, and what can be done to improve outcomes. Neuropharmacology, 120, 11–19. 10.1016/j.neuropharm.2016.03.021

107. Wishart, D. S., Feunang, Y. D., Guo, A. C., Lo, E. J., Marcu, A., Grant, J. R., Sajed, T., Johnson, D., Li, C., Sayeeda, Z., Assempour, N., Iynkkaran, I., Liu, Y., Maciejewski, A., Gale, N., Wilson, A., Chin, L., Cummings, R., Le, D., … Wilson, M. (2017). Drugbank 5.0: A major update to the DrugBank database for 2018. Nucleic Acids Research, 46(D1). 10.1093/nar/gkx1037

108. Davis, A. P., Wiegers, T. C., Wiegers, J., Wyatt, B., Johnson, R. J., Sciaky, D., Barkalow, F., Strong, M., Planchart, A., & Mattingly, C. J. (2023). CTD tetramers: A new online tool that computationally links curated chemicals, genes, phenotypes, and diseases to inform molecular mechanisms for environmental health. Toxicological Sciences, 195(2), 155–168. 10.1093/toxsci/kfad069

109. Azad, A. K., Dinarvand, M., Nematollahi, A., Swift, J., Lutze-Mann, L., & Vafaee, F. (2020). A comprehensive integrated drug similarity resource for in-silico drug repositioning and beyond. Briefings in Bioinformatics, 22(3). 10.1093/bib/bbaa126

110. Szklarczyk, D., Gable, A. L., Lyon, D., Junge, A., Wyder, S., Huerta-Cepas, J., Simonovic, M., Doncheva, N. T., Morris, J. H., Bork, P., Jensen, L. J., & Mering, C. von. (2018). String V11: Protein–protein association networks with increased coverage, supporting functional discovery in genome-wide experimental datasets. Nucleic Acids Research, 47(D1). 10.1093/nar/gky1131

111. Bodenreider O. The Unified Medical Language System (UMLS): integrating biomedical terminology. Nucleic Acids Res. 2004 Jan 1;32(Database issue):D267–70. doi: 10.1093/nar/gkh061

112. Luo, Z., Shi, M., Yang, Z. et al. pyMeSHSim: an integrative python package for biomedical named entity recognition, normalization, and comparison of MeSH terms. BMC Bioinformatics 21, 252 (2020). 10.1186/s12859-020-03583-6

113. Guney, Emre, et al. “Network-Based in Silico Drug Efficacy Screening.” Nature Communications, vol. 7, no. 1, 1 Feb. 2016, 10.1038/ncomms10331.

114. Cheng, Feixiong, et al. “Network-Based Approach to Prediction and Population-Based Validation of in Silico Drug Repurposing.” Nature Communications, vol. 9, no. 1, 12 July 2018, 10.1038/s41467-018-05116-5.

