## Supplemental Document for "From General-Purpose to Disease-Specific Features: Aligning LLM Embeddings on a Disease-Specific Biomedical Knowledge Graph for Drug Repurposing"

**Supplementary Notes 1** | Definitions of F1, AUCROC, and AUPR for Link Prediction

To evaluate link prediction performance of CLEAR and alternative drug repurposing methods, we report the F1 score, Area Under the Receiver Operating Characteristic Curve (AUCROC), and Area Under the Precision-Recall Curve (AUPR). These metrics quantify different aspects of a method's ability to predict the presence or absence of associations between drugs, diseases, and proteins (e.g., drug-disease, drug-protein, disease-protein, or similarity links).

Let a "link" mean a true association between two entities (e.g., drug-disease pair). Then,

- True Positive (**TP**): the model predicts a link between a pair that truly has a link.
- False Positive (**FP**): the model predicts a link between a pair that does not have a true link.
- True Negative (**TN**): the model predicts no link for a pair that truly has no link.
- False Negative (**FN**): the model predicts no link for a pair that truly has a link.

Precision measures the fraction of predicted links that are true links.

$$\mathrm{Pr}\,ecision = \frac{TP}{TP + FP}$$

Recall measures the fraction of true links that are successfully predicted as true links.

$$Recall = \frac{TP}{TP + FN}$$

**F1 score:**

It summarizes *Precision* and *Recall* into a single value by computing their harmonic mean. **Range:** $0 \leq F1\,score \leq 1$ , where **1** indicates perfect *Precision* and *Recall* and **0** indicates that either *Precision* or *Recall* is zero. It is computed as:

$$F1\,score = \frac{\mathrm{Pr}\,ecesion \cdot Recall}{\mathrm{Pr}\,ecesion + Recall}$$

**AUROC:**

It measures how well predicted scores separate true links from non-true links across all classification thresholds. **Range:** $0 \leq AUCROC \leq 1$ , where **1** indicates perfect separability between links and non-links, **0.5** corresponds to random guessing and **0** indicates model is perfectly incorrect (i.e., the model assigns lower scores to all true links than to all non-links). It is defined as the area under the Receiver Operating Characteristic (ROC) curve, which plots the True Positive Rate (TPR) against the False Positive Rate (FPR) where:

$$TPR = \frac{TP}{TP + FN} \; vs. \; FPR = \frac{FP}{FP + TN}$$

AUPR:

It measures the trade-off between *Precision* and *Recall* across thresholds. **Range:** $0 \leq AUPR \leq 1$, where **1** indicates perfect *Precision* and *Recall* across thresholds and **0** indicates complete failure (i.e., the model assigns lower scores to all true links than to all non-links). It is defined as the area under the P*recision-Recall* curve, which plots *Precision* versus *Recall*, where:

$$\Pr e cision = \frac{TP}{TP + FP} \; vs. \; Recall = \frac{TP}{TP + FN}$$

**Supplementary Table 1 |** Statistics of benchmark datasets for drug–disease prediction task.

| Dataset | No. of drugs | No. of diseases | No. of associations |
|---|---|---|---|
| Cdataset | 663 | 409 | 2,352 |
| Fdataset | 593 | 313 | 1,933 |
| Ydataset | 1,478 | 655 | 8,448 |
| LAGCN | 269 | 598 | 18,416 |
| LRSSL | 763 | 681 | 3,051 |

**Supplementary Table 2 |** The **drug-disease association** prediction performances of **SOTA** methods and **CLEAR** on C dataset using 10-fold cross validation.

| *# Unique drugs: 663* | # Unique *diseases: 409* | | # Unique *drug-disease links:2,352* |
|---|---|---|---|
| | ***Metrics*** | | |
| ***Methods*** | F1 | AUPR | AUCROC |
| ITRPCA | 0.6960 | 0.9730 | 0.9654 |
| MLMC | 0.6945 | 0.9747 | 0.9706 |
| OMC | 0.6915 | 0.968 | 0.9586 |
| VDA-GKSBMF | 0.6892 | 0.9628 | 0.9525 |
| BNNR | 0.6889 | 0.9627 | 0.9497 |
| *CLEAR* | **0.959** | **0.988** | **0.9831** |

**Bold** value indicates best performance, and Underlined value indicates second to best performance.

**Supplementary Table 3 |** The **drug-disease association** prediction performances of **SOTA** methods and **CLEAR** on F dataset using 10-fold cross validation.

| *# Unique drugs: 593* | # Unique *diseases: 313* | # Unique *drug-disease links:1,933* | |
|---|---|---|---|
| | | ***Metrics*** | |
| **Methods** | F1 | AUPR | AUCROC |
| HINGRL | <u>0.8789</u> | 0.9515 | 0.9432 |
| ITRPCA | 0.6896 | 0.963 | 0.9529 |
| MLMC | 0.6895 | <u>0.9652</u> | <u>0.9594</u> |
| OMC | 0.685 | 0.9559 | 0.9436 |
| DRRS | 0.6809 | 0.9497 | 0.9343 |
| *CLEAR* | **0.9543** | **0. 9893** | **0.9858** |

**Bold** value indicates best performance, and <u>Underlined</u> value indicates second to best performance.

**Supplementary Table 4 |** The **drug-disease association** prediction performances of **SOTA** methods and **CLEAR** on Y dataset using 10-fold cross validation.

| *# Unique drugs: 1,478* | # Unique *diseases: 655* | # Unique *drug-disease links:8,448* | |
|---|---|---|---|
| | | ***Metrics*** | |
| **Methods** | F1 | AUPR | AUCROC |
| VDA-GKSBMF | <u>0.6981</u> | <u>0.9751</u> | <u>0.9699</u> |
| DRRS | 0.697 | 0.9676 | 0.9593 |
| ITRPCA | 0.6955 | 0.9712 | 0.9619 |
| DRPADC | 0.6944 | 0.9689 | 0.9618 |
| HGIMC | 0.6923 | 0.9662 | 0.9574 |
| *CLEAR* | **0.9219** | **0.9776** | **0.9741** |

**Bold** value indicates best performance, and <u>Underlined</u> value indicates second to best performance.

**Supplementary Table 5 |** The **drug-disease association** prediction performances of **SOTA** methods and **CLEAR** on LAGCN dataset using 10-fold cross validation.

*# Unique drugs: 269* | # Unique *diseases: 598* | # Unique *drug-disease links:18,416*

| ***Methods*** | ***Metrics*** | | |
|---|---|---|---|
| | F1 | AUPR | AUCROC |
| SCMFDD | 0.4316 | 0.828 | **0.9665** |
| GROBMC | 0.6917 | 0.6187 | 0.9493 |
| HINGRL | 0.815 | 0.8768 | 0.8853 |
| LAGCN | 0.2355 | 0.2618 | 0.8783 |
| DRHGCN | 0.5354 | 0.5559 | 0.8756 |
| *CLEAR* | **0.8315** | **0.8887** | 0.9057 |

**Bold** value indicates best performance, and Underlined value indicates second to best performance.

**Supplementary Table 6 |** The **drug-disease association** prediction performances of **SOTA** methods and **CLEAR** on LRSSL dataset using 10-fold cross validation.

*# Unique drugs: 763* | # Unique *diseases: 681* | # Unique *drug-disease links:3,051*

| **Methods** | ***Metrics*** | | |
|---|---|---|---|
| | F1 | AUPR | AUCROC |
| GROBMC | 0.6134 | 0.588 | 0.9594 |
| DRHGCN | 0.499 | 0.448 | 0.9583 |
| DRIMC | 0.0061 | 0.162 | 0.9533 |
| OMC | 0.6853 | 0.9545 | 0.9437 |
| DDAPRED | 0.4973 | 0.4672 | 0.9381 |
| *CLEAR* | **0.9863** | **0.9973** | **0.9862** |

**Bold** value indicates best performance, and Underlined value indicates second to best performance.

**Supplementary Table 7 |** The **drug-disease association** prediction performances of **SOTA** methods and **CLEAR** on ADRD dataset using 10-fold cross validation.

| *# Unique drugs: 269* | # Unique *diseases: 598* | # Unique *drug-disease links:18,416* | |
|---|---|---|---|
| | ***Metrics*** | | |
| ***Methods*** | F1 | AUPR | AUCROC |
| HINGRL | <u>0.815</u> | <u>0.865</u> | 0.8990 |
| GROBMC | 0.0004 | 0.04 | 0.465 |
| SCMFDD | 0.0476 | 0.0509 | 0.8968 |
| ITRPCA | 0.1505 | 0.0683 | 0.938 |
| VDA-GKSBMF | 0.0004 | 0.2144 | <u>0.9709</u> |
| *CLEAR* | **0.9887** | **0.9964** | **0.9969** |

**Bold** value indicates best performance, and <u>Underlined</u> value indicates second to best performance.

**Supplementary Table 8 |** Top 20 Significantly enriched **Biological Process (BP)** Gene Ontology (GO) terms for the therapeutic target proteins of **Dextromethorphan**. (False Discovery Rate (FDR) ≤ 0.05)

| *GO terms ID* | GO term description | Enrichment FDR | #Therapeutic Proteins | #Proteins in GO term | Fold Enrichment |
|---|---|---|---|---|---|
| *GO:0045730* | Respiratory burst | 2.16E-12 | 7 | 37 | 206.135 |
| *GO:0007271* | Synaptic transmission cholinergic | 1.18E-10 | 6 | 31 | 210.885 |
| *GO:0006810* | Transport | 1.46E-10 | 20 | 4823 | 4.518 |
| *GO:0060078* | Reg. Of postsynaptic membrane potential | 1.46E-10 | 8 | 145 | 60.114 |
| *GO:0051234* | Establishment of localization | 2.26E-10 | 20 | 4987 | 4.370 |
| *GO:0034220* | Ion transmembrane transport | 6.04E-10 | 13 | 1218 | 11.629 |
| *GO:0060079* | Excitatory postsynaptic potential | 6.04E-10 | 7 | 102 | 74.775 |
| *GO:0099565* | Chemical synaptic transmission postsynaptic | 7.53E-10 | 7 | 109 | 69.972 |
| *GO:0006811* | Ion transport | 1.15E-09 | 14 | 1677 | 9.096 |
| *GO:0035094* | Response to nicotine | 1.15E-09 | 6 | 57 | 114.692 |
| *GO:0007268* | Chemical synaptic transmission | 1.39E-09 | 11 | 764 | 15.688 |
| *GO:0098916* | Anterograde trans-synaptic signaling | 1.39E-09 | 11 | 764 | 15.688 |
| *GO:0099537* | Trans-synaptic signaling | 1.45E-09 | 11 | 773 | 15.505 |
| *GO:0099536* | Synaptic signaling | 2.23E-09 | 11 | 810 | 14.797 |
| *GO:0030534* | Adult behavior | 3.88E-09 | 7 | 150 | 50.847 |
| *GO:0035095* | Behavioral response to nicotine | 4.92E-09 | 4 | 9 | 484.254 |
| *GO:0042391* | Reg. Of membrane potential | 5.65E-09 | 9 | 442 | 22.186 |
| *GO:0051046* | Reg. Of secretion | 5.65E-09 | 10 | 650 | 16.763 |
| *GO:0055085* | Transmembrane transport | 1.03E-08 | 13 | 1653 | 8.569 |
| *GO:0038003* | G protein-coupled opioid receptor signaling pathway | 2.12E-08 | 4 | 13 | 335.253 |

**Supplementary Table 9 |** Top 20 Significantly enriched **Molecular Function (MF)** Gene Ontology (GO) terms for the therapeutic target proteins of **Dextromethorphan**. (False Discovery Rate (FDR) ≤ 0.05)

| *GO terms ID* | GO term description | Enrichment FDR | #Therapeutic Proteins | #Proteins in GO term | Fold Enrichment |
|---|---|---|---|---|---|
| *GO:0022848* | Acetylcholine-gated cation-selective channel activity | 1.35E-12 | 6 | 20 | 326.871 |
| *GO:1904315* | Transmitter-gated ion channel activity involved in reg. Of postsynaptic membrane potentia | 1.35E-12 | 7 | 50 | 152.540 |
| *GO:0015464* | Acetylcholine receptor activity | 1.41E-12 | 6 | 24 | 272.393 |
| *GO:0022824* | Transmitter-gated ion channel activity | 2.48E-12 | 7 | 63 | 121.063 |
| *GO:0022835* | Transmitter-gated channel activity | 2.48E-12 | 7 | 63 | 121.063 |
| *GO:0098960* | Postsynaptic neurotransmitter receptor activity | 4.13E-12 | 7 | 69 | 110.536 |
| *GO:0005231* | Excitatory extracellular ligand-gated ion channel activity | 5.76E-12 | 6 | 33 | 198.104 |
| *GO:0005230* | Extracellular ligand-gated ion channel activity | 7.13E-12 | 7 | 77 | 99.052 |
| *GO:0042165* | Neurotransmitter binding | 6.96E-11 | 5 | 19 | 286.729 |
| *GO:0030594* | Neurotransmitter receptor activity | 9.18E-11 | 7 | 113 | 67.496 |
| *GO:0099094* | Ligand-gated cation channel activity | 1.29E-10 | 7 | 120 | 63.558 |
| *GO:0022836* | Gated channel activity | 2.23E-10 | 9 | 374 | 26.220 |
| *GO:0022834* | Ligand-gated channel activity | 5.27E-10 | 7 | 151 | 50.510 |
| *GO:0004985* | G protein-coupled opioid receptor activity | 6.36E-10 | 4 | 9 | 484.254 |

| | | | | | |
|---|---|---|---|---|---|
| *GO:0022890* | Inorganic cation transmembrane transporter activity | 6.36E-10 | 10 | 637 | 17.105 |
| *GO:0015075* | Ion transmembrane transporter activity | 8.95E-10 | 11 | 922 | 12.999 |
| *GO:0016175* | Superoxide-generating NAD(P)H oxidase activity | 8.95E-10 | 4 | 10 | 435.829 |
| *GO:0042166* | Acetylcholine binding | 2.75E-09 | 4 | 13 | 335.253 |
| *GO:0015267* | Channel activity | 2.87E-09 | 9 | 530 | 18.502 |
| *GO:0022803* | Passive transmembrane transporter activity | 2.87E-09 | 9 | 532 | 18.433 |

**Supplementary Table 10 |** Top 20 Significantly enriched **Cellular Component (CC)** Gene Ontology (GO) terms for the therapeutic target proteins of ***Dextromethorphan***. (False Discovery Rate (FDR) ≤ 0.05)

| ***GO terms ID*** | **GO term description** | **Enrichment FDR** | **#Therapeutic Proteins** | **#Proteins in GO term** | **Fold Enrichment** |
|---|---|---|---|---|---|
| *GO:0005887* | Integral component of plasma membrane | 7.57E-19 | 20 | 1894 | 11.506 |
| *GO:0031226* | Intrinsic component of plasma membrane | 9.02E-19 | 20 | 1978 | 11.017 |
| *GO:0043005* | Neuron projection | 1.51E-17 | 18 | 1440 | 13.620 |
| *GO:0043020* | NADPH oxidase complex | 7.65E-17 | 7 | 16 | 476.688 |
| *GO:0097060* | Synaptic membrane | 7.65E-17 | 13 | 395 | 35.859 |
| *GO:0045211* | Postsynaptic membrane | 1.05E-16 | 12 | 288 | 45.399 |
| *GO:0120025* | Plasma membrane bounded cell projection | 7.75E-16 | 19 | 2360 | 8.772 |
| *GO:0042995* | Cell projection | 1.69E-15 | 19 | 2477 | 8.358 |
| *GO:0098797* | Plasma membrane protein complex | 1.98E-15 | 14 | 730 | 20.896 |
| *GO:0098794* | Postsynapse | 2.85E-14 | 13 | 664 | 21.332 |
| *GO:0005892* | Acetylcholine-gated channel complex | 8.59E-14 | 6 | 18 | 363.190 |
| *GO:0098590* | Plasma membrane region | 1.69E-13 | 15 | 1332 | 12.270 |
| *GO:0045202* | Synapse | 4.67E-13 | 15 | 1435 | 11.389 |
| *GO:0030054* | Cell junction | 5.32E-13 | 17 | 2293 | 8.078 |
| *GO:0098796* | Membrane protein complex | 1.55E-11 | 14 | 1452 | 10.506 |
| *GO:0036477* | Somatodendritic compartment | 2.71E-11 | 12 | 888 | 14.724 |
| *GO:0043025* | Neuronal cell body | 9.20E-11 | 10 | 513 | 21.239 |
| *GO:0044297* | Cell body | 3.33E-10 | 10 | 588 | 18.530 |
| *GO:1990204* | Oxidoreductase complex | 5.80E-10 | 7 | 154 | 49.526 |
| *GO:0098802* | Plasma membrane signaling receptor complex | 3.46E-09 | 7 | 200 | 38.135 |

**Supplementary Table 11 |** The **drug-disease association** prediction performances of **baseline Machine Learning** using LLM embeddings vs. using CLEAR embeddings on ADRD dataset using 10-fold cross validation.

| Model | F1 | | AUPR | | AUCROC | |
|---|---|---|---|---|---|---|
| | *LLM embedding* | *CLEAR embedding* | *LLM embedding* | *CLEAR embedding* | *LLM embedding* | *CLEAR embedding* |
| RF | 0.851 (0.003) | **0.860 (0.005)** | 0.929 (0.003) | **0.937 (0.004)** | 0.925 (0.002) | **0.933 (0.003)** |
| SVM | 0.818 (0.005) | **0.831 (0.004)** | 0.892 (0.005) | **0.904 (0.002)** | 0.886 (0.007) | **0.898 (0.004)** |
| MLP | 0.850 (0.006) | **0.906 (0.003)** | 0.922 (0.006) | **0.967 (0.003)** | 0.912 (0.009) | **0.963 (0.004)** |
| XGBoost | 0.860 (0.002) | **0.897 (0.005)** | 0.930 (0.002) | **0.959 (0.004)** | 0.918 (0.004) | **0.956 (0.005)** |

Values outside the bracket indicate **mean score** and values inside the brackets indicate **standard deviation of the score** across 10 folds. For each metric and ML method, higher performance value is shown in ***bold.***

**Supplementary Table 12 |** The **drug-disease association** prediction performances of **ablated variants** of CLEAR on ADRD dataset using 10-fold cross validation.

| | ***Model*** | **F1** | **AUPR** | **AUCROC** |
|---|---|---|---|---|
| Computational modules ablation | *Variant 1* | 0.8640 ± 0.0230 | 0.9526 ± 0.0104 | <u>0.944</u> ± <u>0.0081</u> |
| | *Variant 2* | 0.8758 ± 0.002 | 0.9182 ± 0.002 | 0.9044 ± 0.004 |
| | *Variant 3* | <u>0.9546</u> ± <u>0.0110</u> | <u>0.9700</u> ± <u>0.001</u> | 0.9353 ± 0.009 |
| | *Variant 4* | 0.9467 ± 0.002 | 0.9753 ± 0.002 | 0.9553 ± 0.004 |
| Data ablati on | *Variant 5* | 0.75532 ± 0.0391 | 0.9213 ± 0.0347 | 0.8457 ± 0.0378 |
| | *Variant 6* | 0.8703 ± 0.0357 | 0.9325 ± 0.0296 | 0.9369 ± 0.0329 |
| | *CLEAR (Proposed method)* | **0. 9887 ± 0.001** | **0. 9964 ±0.003** | **0. 9969 ± 0.002** |

**Bold** value indicates best performance, and <u>Underlined</u> value indicates second to best performance

**Variant 1**: CLEAR trained without GAT and MHSA
**Variant 2**: CLEAR trained without MHSA
**Variant 3**: CLEAR trained using non-weighted loss function
**Variant 4**: CLEAR trained using randomly selected negative edges

**Variant 5**: CLEAR trained using randomly initialized node features
**Variant 6**: CLEAR trained using Drug and Disease related data only by removing all Protein nodes from training.

**CLEAR**: CLEAR with all modules and data types.

**Supplementary Table 13** | Downstream drug repurposing rankings using CLEAR trained with sparse data. The table lists the top five candidate drugs predicted by CLEAR for Alzheimer's Disease (AD), Parkinson's Disease Dementia (PDD), and Lewy Body Dementia (LBD) when trained with *25% drug-disease links*, *0% LLM-derived node embeddings*, and varying proportions of *other links (0%, 1%, 5%, and 10%)*. Rankings remain largely the same across diseases and other links settings, indicating limited disease-specific differentiation under sparse training. Note: Drug names shared across two or more diseases are underlined, while those shared across two or more training settings are **bold**. Drugs that are both **bold and underlined** are shared across multiple diseases and training settings.

| *LLM embedding* | | 0% | | | |
|---|---|---|---|---|---|
| *Drug-Disease links* | | 25% | | | |
| *Other links* | | 0% | 1% | 5% | 10% |
| | **Rank** | **Drug Name** | **Drug Name** | **Drug Name** | **Drug Name** |
| *Alzheimer Disease* | 1 | Viomycin | Disulfiram | Iopodic acid | Artenimol |
| | 2 | Casirivimab | Mezlocillin | Sertaconazole | Indacaterol |
| | 3 | Rituximab | Nitric Oxide | Peginterferon beta-1a | Zinc acetate |
| | 4 | Ombitasvir | Potassium | Calcium carbimide | Zinc |
| | 5 | Tezepelumab | Triamcinolone | Axicabtagene ciloleucel | Brompheniramine |
| *Parkinson Disease Dementia* | 1 | Viomycin | Disulfiram | Iopodic acid | Indacaterol |
| | 2 | Ombitasvir | Mezlocillin | Peginterferon beta-1a | Brompheniramine |
| | 3 | Rituximab | Nitric Oxide | Sertaconazole | Ifosfamide |
| | 4 | Casirivimab | Potassium | Calcium carbimide | Midodrine |
| | 5 | Galsulfase | Triamcinolone | Axicabtagene ciloleucel | Quinine |
| *Lewy Body Dementia* | 1 | Viomycin | **Amitriptyline** | Iopodic acid | Fostamatinib |
| | 2 | Casirivimab | Triamcinolone | Peginterferon beta-1a | **Amitriptyline** |
| | 3 | Rituximab | Copper | Sertaconazole | Zinc acetate |
| | 4 | Ombitasvir | Nandrolone decanoate | Calcium carbimide | Zinc |
| | 5 | Tezepelumab | Esketamine | Axicabtagene ciloleucel | Zinc chloride |

**Supplementary Table 14 |** Downstream drug repurposing rankings using CLEAR trained with sparse data. The table lists the top five candidate drugs predicted by CLEAR for Alzheimer's Disease (AD), Parkinson's Disease Dementia (PDD), and Lewy Body Dementia (LBD) when trained with *25% drug-disease links*, *100% LLM-derived node embeddings*, and varying proportions of *other links (0%, 1%, 5%, and 10%)*. Rankings remain largely the same across diseases and other links settings, indicating limited disease-specific differentiation under sparse training. Note: Drug names shared across two or more diseases are underlined, while those shared across two or more training settings are **bold**. Drugs that are both **bold and underlined** are shared across multiple diseases and training settings.

| *LLM embedding* | | 100% | | | |
|---|---|---|---|---|---|
| *Drug-Disease links* | | 25% | | | |
| *Other links* | | 0% | 1% | 5% | 10% |
| | **Rank** | **Drug Name** | **Drug Name** | **Drug Name** | **Drug Name** |
| *Alzheimer Disease* | 1 | **Dequalinium** | **Dequalinium** | Docetaxel | **Dequalinium** |
| | 2 | **Diphemanil** | **Oxymetazoline** | **Zinc chloride** | **Tetanus immune globulin** |
| | 3 | **Oxymetazoline** | **Diphemanil** | Isotretinoin | **Diphemanil** |
| | 4 | **Droxidopa** | **Droxidopa** | Norgestimate | **Oxymetazoline** |
| | 5 | **Ephedrine** | **Ephedrine** | Cyclophosphamide | **Droxidopa** |
| *Parkinson Disease Dementia* | 1 | **Tetanus immune globulin** | **Procaine** | Amphetamine | **Metamfetamine** |
| | 2 | **Metamfetamine** | **Haloperidol** | **Procaine** | **Amoxapine** |
| | 3 | **Amoxapine** | **Mianserin** | **Haloperidol** | **Mianserin** |
| | 4 | **Mianserin** | **Trimipramine** | Aripiprazole | **Loxapine** |
| | 5 | Triflupromazine | **Loxapine** | **Zinc chloride** | **Trimipramine** |
| *Lewy Body Dementia* | 1 | **Dequalinium** | **Tetanus immune globulin** | Docetaxel | **Dequalinium** |
| | 2 | **Tetanus immune globulin** | **Oxymetazoline** | **Zinc chloride** | **Tetanus immune globulin** |
| | 3 | **Diphemanil** | **Droxidopa** | Isotretinoin | **Diphemanil** |
| | 4 | **Oxymetazoline** | **Zinc chloride** | Norgestimate | **Oxymetazoline** |
| | 5 | Droxidopa | **Diphemanil** | Cyclophosphamide | **Droxidopa** |

**Supplementary Figure 1 |** The **drug-disease association**, **drug-protein association** and **disease-protein association** prediction performances of **CLEAR** on benchmark datasets and ADRD datasets.

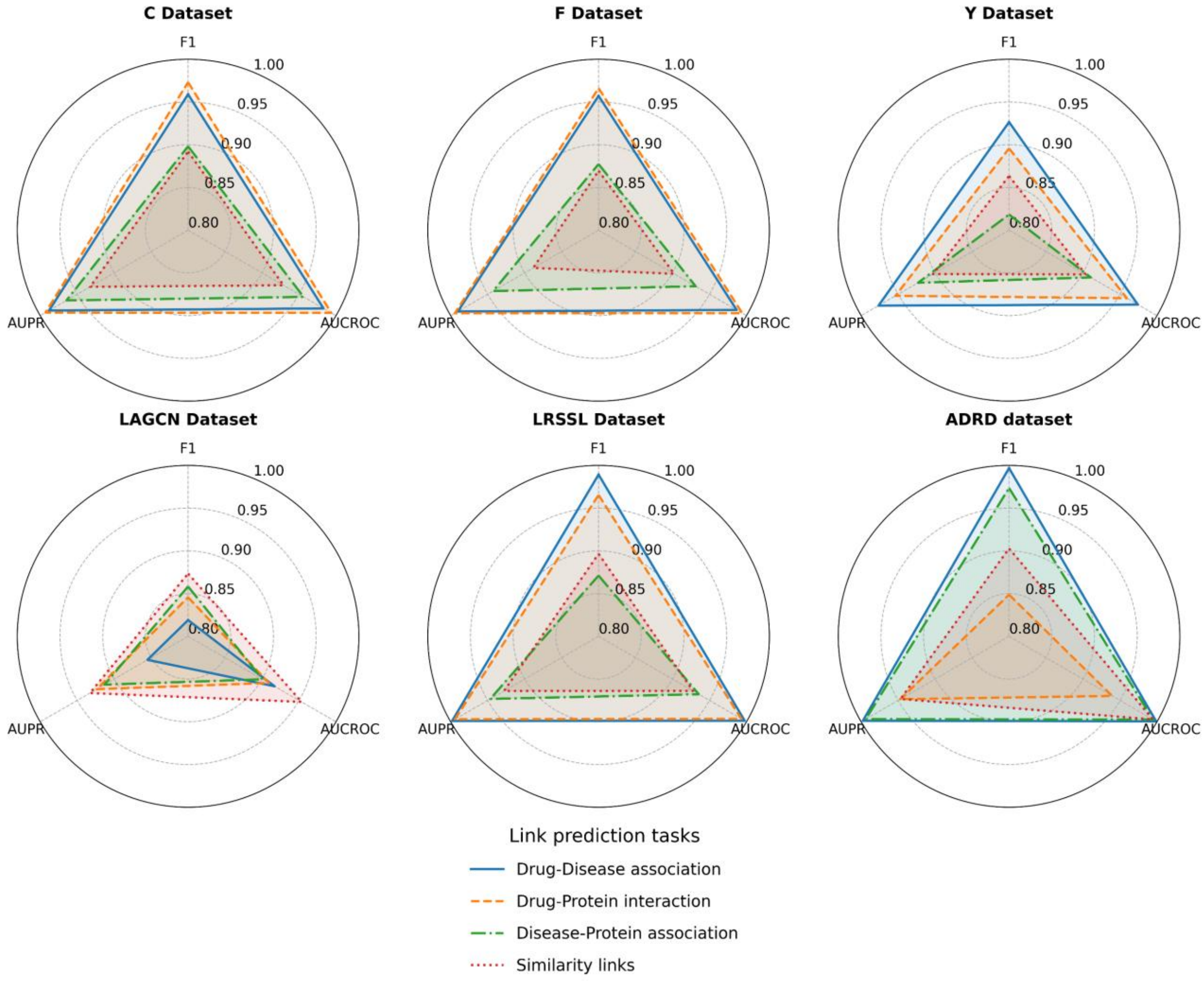

**Supplementary Figure 2 |** Plot showing overlap of drug–disease links across benchmark datasets. Vertical bars indicate the number of shared links for each dataset combination, as marked by the connected dots in the matrix below. Horizontal bars show the total number of links in each dataset.

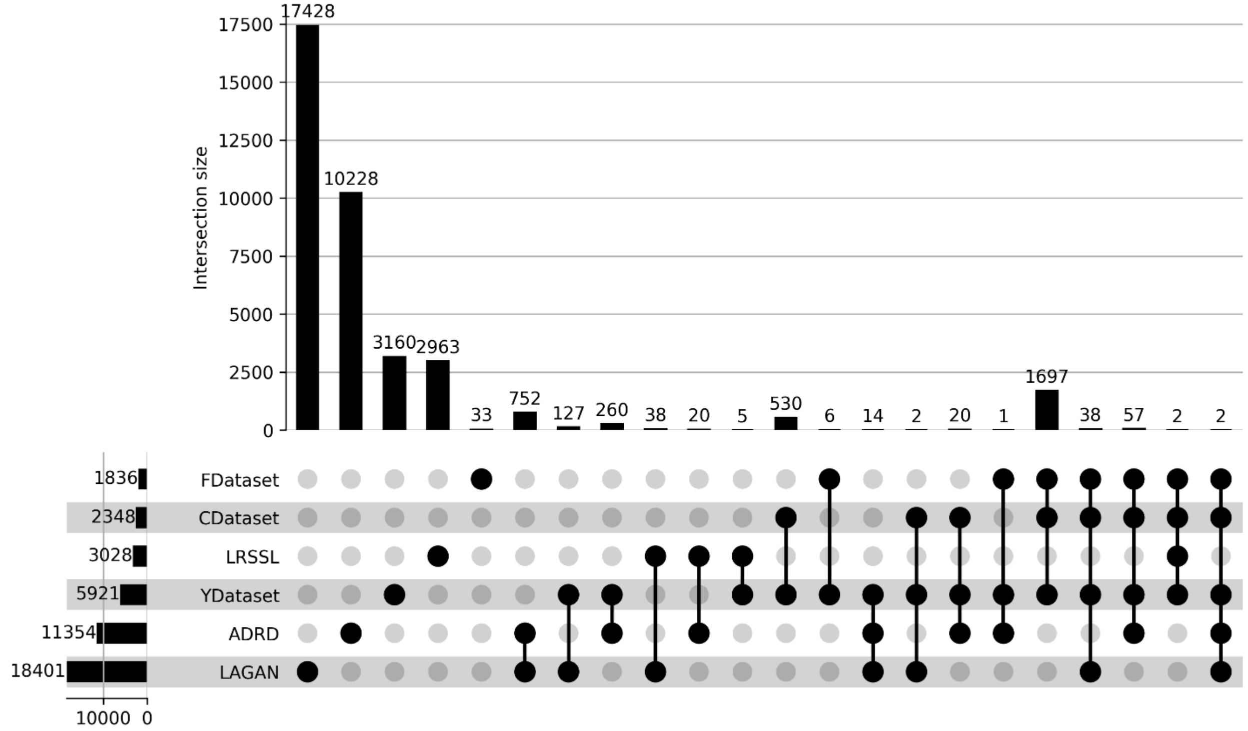

**Supplementary Figure 3 |** Performance of CLEAR on the *drug-disease link* prediction task across different levels of training data sparsity. a. F1 score, b. AUROC, and c. AUPR across varying percentages of *LLM node embeddings* used during CLEAR training. Marker color indicates the percentage of *drug-disease links* used during CLEAR training (0%, 1%, 5%, 10%, and 25%), while marker shape denotes the percentage of *other links* used during CLEAR training (0%, 1%, 5%, and 10%). Across all metrics, performance saturates once 25% of *drug-disease links* are available, with only marginal gains from increasing the percentage of *other links* or *node embedding*, highlighting that strong predictive performance can be achieved under sparse supervision. Notably, only three marker shapes (square, triangle, and diamond) are shown: when 0% of drug-disease links are used, 1% of other links are included to avoid an edgeless graph, as CLEAR requires at least one link type to propagate information.

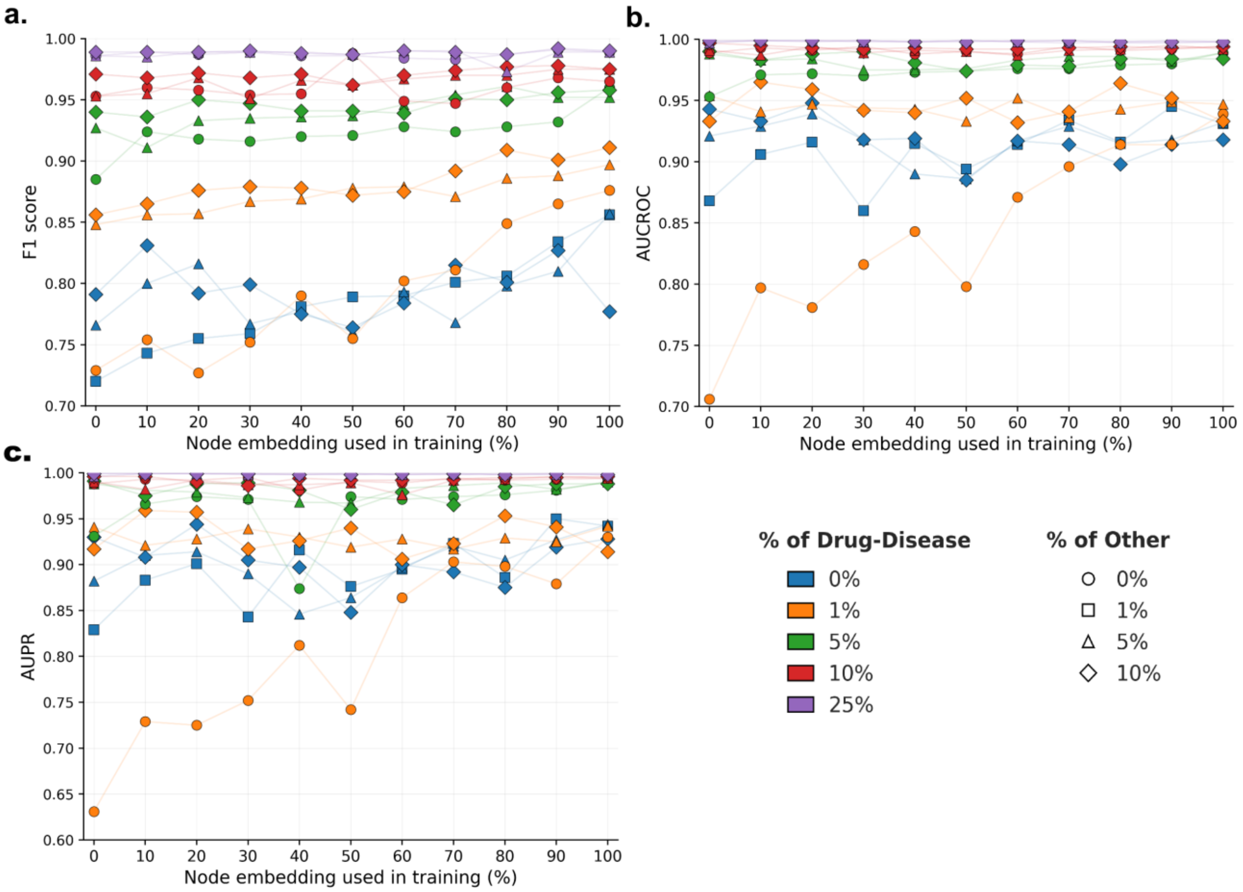

**Supplementary Figure 4** | Computational resource usage of the CLEAR framework. The figure illustrates the key computational metrics for the CLEAR model, including (a) the total number of trainable parameters, (b) peak GPU memory usage during training, and (c) total compute time required for a full training run.

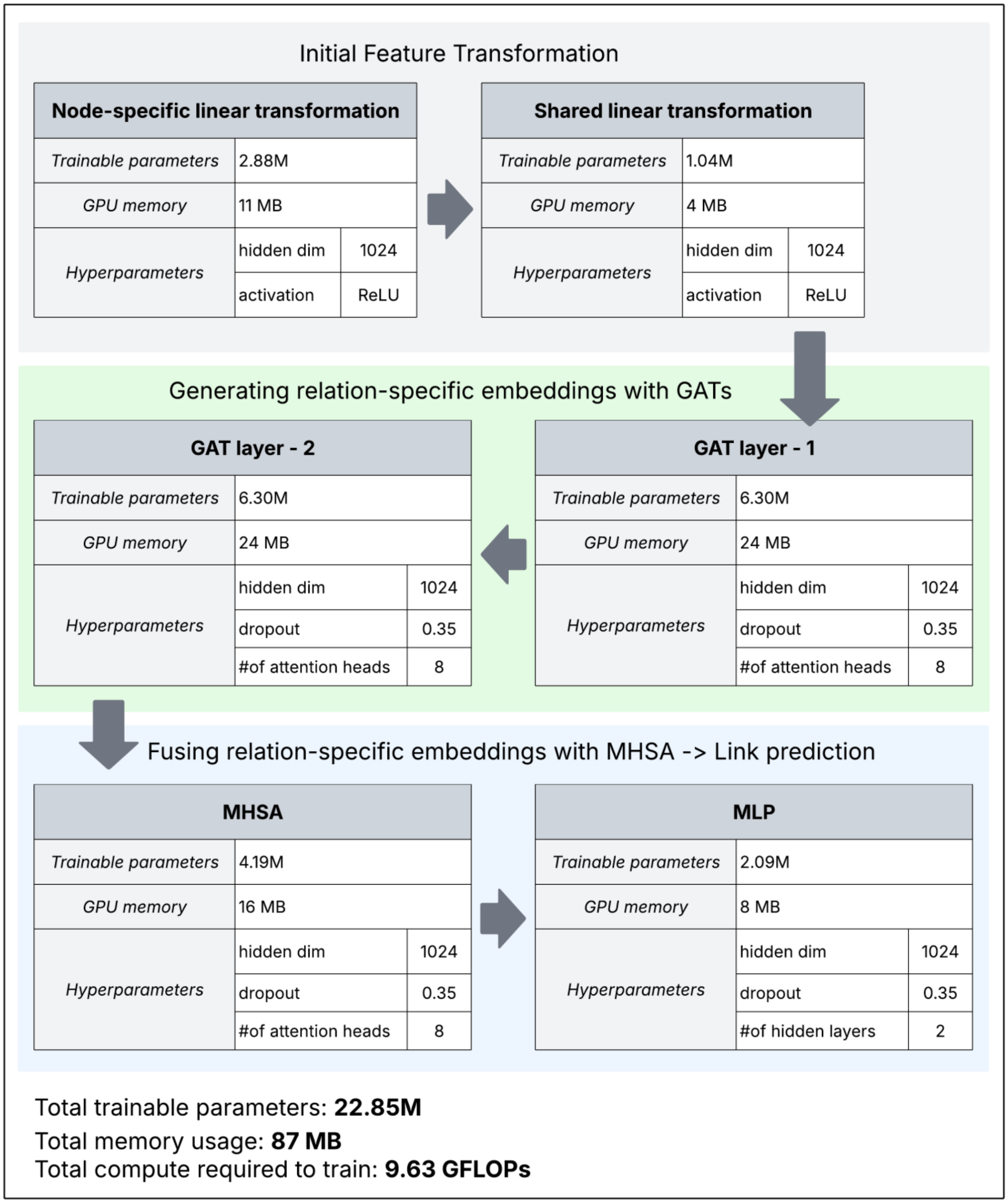

**Supplementary Algorithm 1 |** Negative node pair sampling

**Input**: $dict_{neighbors}$ : $v_i \rightarrow 3\ hop\ neighbors\ of\ v_i$ , knowledge graph $G_{RDT}(V, E)$

**Output**: Positive node pairs $P$ and negative node pairs $N$

Decompose $G_{RDT}(V, E)$ into subgraphs:

$G = \{G_R, G_D, G_T, G_{R-D}, G_{R-T}, G_{D-T}\}$

**for each** $G_i(V, E) \in G$ **do**:

$P_i \leftarrow \{(u, v) | (u, v) \in E(G_i)\}$

Compute positive pair degree: $\deg(u, v) = \deg(u) + \deg(v)$

$p_i(d) \leftarrow freq_{(u,v) \in P_i}(\deg(u, v) = d)$

$S_i \leftarrow source\ node\ set\ of\ G_i,$

$D_i \leftarrow destination\ node\ set\ of\ G_i,$

$N_i \leftarrow \emptyset$

**while** $|N_i| < |P_i|$ **do**:

$d_s \sim p_i(d)$

$u \sim Uniform(s_i)$

#3-hop exclusion, prevents false negatives by skipping nearby nodes

$C(u) = \{v \in D_i \mid v \notin dict_{neighbors}[u], v \neq u\}$

**repeat up to $n$ times:**

$v \sim Uniform\ (C(u))$

**if** $(u, v) \notin E(G_i)$ **and** $|\deg(u, v) - d_s| < \varepsilon$ **then**

#degree matching, creates "hard" negatives pairs that is structurally similar to positives pairs

$N_i \leftarrow N_i \cup \{(u, v)\}$

**break**

**end if**

**end repeat**

**end while**

### subgraph balance, ensures equal representation of all relation types

$P \leftarrow P \cup P_i, N \leftarrow N \cup N_i$

**return** $\{P, N\}$

**Supplementary Algorithm 2 |** Relation-specific feature update

**Input**: Knowledge graph $G_{R\text{-}D\text{-}T}(V,E)$, LLM-based drug features $\{x_R \in \mathbb{R}^{v*r}, \forall v \in V_R\}$, LLM-based disease features $\{x_D \in \mathbb{R}^{v*d}, \forall v \in V_D\}$, LLM-based protein features $\{x_T \in \mathbb{R}^{v*t}, \forall v \in V_T\}$, edge level node embedding aggregator function AGGREGATE

**Output**: drug node embedding $\{Z_R \in \mathbb{R}^{v*d}, \forall v \in V_R\}$, disease node embedding $\{Z_D \in \mathbb{R}^{v*d}, \forall v \in V_R\}$, protein node embedding $\{Z_T \in \mathbb{R}^{v*d}, \forall v \in V_R\}$

***Graph decomposition***

$A_{R\text{-}sim}$, $A_{D\text{-}sim}$, $A_{T\text{-}sim} \leftarrow G_{R\text{-}D\text{-}T}(V, E)$, similarity adjacency matrix

$A_{R\text{-}D}$, $A_{R\text{-}T}$, $A_{D\text{-}T} \leftarrow G_{R\text{-}D\text{-}T}(V, E)$, bipartite adjacency matrix

where $G_{R\text{-}D\text{-}T}(V, E)$ is a drug-disease-protein knowledge graph

***Feature transformation***

Drug input feature $\{x_R \in \mathbb{R}^{v*r}, \ \forall v \in V_R\}$

Disease input feature $\{x_D \in \mathbb{R}^{v*d}, \ \forall v \in V_D\}$

Protein input feature $\{x_T \in \mathbb{R}^{v*t}, \ \forall v \in V_T\}$,

*Feature-specific transformation*

$\overline{x_R} \leftarrow x_R W_R$, where $W_R \in \mathbb{R}^{r*f}$

$\overline{x_D} \leftarrow x_D W_D$, where $W_D \in \mathbb{R}^{d*f}$

$\overline{x_T} \leftarrow x_T W_T$, where $W_T \in \mathbb{R}^{t*f}$

*Shared feature transformation*

$\overline{\overline{x_R}} \leftarrow \overline{x_R} \ W_{shared}$

$\overline{\overline{x_D}} \leftarrow \overline{x_D} \ W_{shared}$

$\overline{\overline{x_T}} \leftarrow \overline{x_T} \ W_{shared}$, where $W_{shared} \in \mathbb{R}^{f*g}$

***Feature update through relation specific GATs (similarity relationships)***

$H^{(2)}_{R-sim}$ ← GAT ($A_{R\text{-}sim}$, $f$(GAT ($A_{R\text{-}sim}$, $\overline{\overline{x_R}}$ , $W_R^{(1)}$), $W_R^{(2)}$)

$H^{(2)}_{D-sim}$ ← GAT ($A_{D\text{-}sim}$, $f$(GAT ($A_{D\text{-}sim}$, $\overline{\overline{x_R}}$ , $W_D^{(1)}$), $W_D^{(2)}$)

$H^{(2)}_{T-sim}$ ← GAT ($A_{T\text{-}sim}$, $f$(GAT ($A_{T\text{-}sim}$, $\overline{\overline{x_R}}$ , $W_T^{(1)}$), $W_T^{(2)}$), where $f$ is ReLU activation function

***Feature update through relation specific GATs (bipartite relationships)***

$\overline{\overline{x_{R+D}}} \leftarrow \overline{\overline{x_R}} \,||\, \overline{\overline{x_D}}$ , where $\overline{\overline{x_{R+D}}} \in \mathbb{R}^{(r+d) * g}$

$\overline{\overline{x_{R+T}}} \leftarrow \overline{\overline{x_R}} \,||\, \overline{\overline{x_T}}$ , where $\overline{\overline{x_{R+T}}} \in \mathbb{R}^{(r+t) * g}$

$\overline{\overline{x_{D+T}}} \leftarrow \overline{\overline{x_D}} \,||\, \overline{\overline{x_T}}$ , where $\overline{\overline{x_{D+T}}} \in \mathbb{R}^{(d+t) * g}$

$H^{(2)}_{R-D}$ ← GAT ($A_{R\text{-}D}$, $f$(GAT ($A_{R\text{-}D}$, $\overline{\overline{x_R}}$ , $W^{(1)}_{R-D}$), $W^{(2)}_{R-D}$)

$H^{(2)}_{R-T}$ ← GAT ($A_{R\text{-}T}$, $f$(GAT ($A_{R\text{-}T}$, $\overline{\overline{x_R}}$ , $W^{(1)}_{R-T}$), $W^{(2)}_{R-T}$)

$H^{(2)}_{D-T}$ ← GAT ($A_{D\text{-}T}$, $f$(GAT ($A_{D\text{-}T}$, $\overline{\overline{x_R}}$ , $W^{(1)}_{D-T}$), $W^{(2)}_{D-T}$), where $f$ is ReLU activation function

***Multi-relation feature fusion through MHSA***

$Z^R_{CLEAR} \leftarrow fuse\ (H^{(2)}_{R-sim}\ , H^{(1)}_{R-D}\ , H^{(2)}_{R-T})$, where $Z^R_{CLEAR} \in \mathbb{R}^{r * h}$

$Z^D_{CLEAR} \leftarrow fuse\ (H^{(2)}_{R-sim}\ (H^{(2)}_{D-sim}\ , H^{(1)}_{R-D}\ , H^{(2)}_{D-T})$, where $Z^D_{CLEAR} \in \mathbb{R}^{d * h}$

$Z^T_{CLEAR} \leftarrow fuse\ (H^{(2)}_{R-sim}\ (H^{(2)}_{T-sim}\ , H^{(1)}_{R-T}\ , H^{(2)}_{D-T})$, where $Z^T_{CLEAR} \in \mathbb{R}^{t * h}$

---

**Supplementary Algorithm 3 |** Multi-relation feature fusion

**Input**: single updated feature matrix based on similarity $GAT_{sim}$, $H_{sim}^{(2)} \in \mathbb{R}^{\mathrm{r,d,t*h}}$ ;
Two of the updated feature matrices based on bipartite $GAT_{bi\_1}$ and $GAT_{bi\_2}$ ,
$H_{bi\_1}^{(2)}$ $and$ $H_{bi\_2}^{(2)} \in \mathbb{R}^{\mathrm{r,d,t*h}}$

**Output**: CLEAR embeddings $Z_{CLEAR} \in \mathbb{R}^{\mathrm{r,d,t*h}}$

***Multi-head queries, keys and values***

X ← Stack ($H_{sim}^{(2)}$ , $H_{bi\_1}^{(2)}$ , $H_{bi\_2}^{(2)}$ )

$Query_i \leftarrow X\, W_i^Q$ ,

$Key_i \leftarrow X\, W_i^K$ ,

$Value_i \leftarrow X\, W_i^V$, where $W_i^Q$, $W_i^D$ and $W_i^T \in \mathbb{R}^{\mathrm{r,d,t*d}}$ are learnable weight matrices for $Query$, $Key$ and $Value$, respectively for $i^{th}$ attention head.

***Compute attention score per head***

$Attention_{\,i}\,(Query_i\,, Key_i\,, Value_i\,) = \sigma\left(\frac{Query_i * {Key_i}^T}{\sqrt{d_{Key_i}}}\right) Value_i$, where $d_{Key}$ is the dimension of the $Key_i$ vectors.

***Combine multi-head attention scores to generate CLEAR embedding***

$Z_{CLEAR} \leftarrow Concat(Attention_{\,1}, \dots Attention_{\,h})\, W_O$ , where $W_O \in \mathbb{R}^{\mathrm{r,d,t*h}}$ is a final weight matrix to project the concatenated output back to embedding dimension.

**Supplementary Algorithm 4 |** Link prediction

**Input**: Drug, disease and protein's CLEAR embedding, $Z^{R}_{CLEAR}$, $Z^{D}_{CLEAR}$and $Z^{T}_{CLEAR}$

**Output:** Binary link prediction score between source node and destination node.

***Bipartite link prediction***

$Z^{RD}_{CLEAR} \leftarrow Z^{R}_{CLEAR} \,||\, Z^{D}_{CLEAR} \in \mathbb{R}^{2*\mathrm{h}}$

$Z^{RT}_{CLEAR} \leftarrow Z^{R}_{CLEAR} \,||\, Z^{T}_{CLEAR} \in \mathbb{R}^{2*\mathrm{h}}$

$Z^{DT}_{CLEAR} \leftarrow Z^{D}_{CLEAR} \,||\, Z^{T}_{CLEAR} \in \mathbb{R}^{2*\mathrm{h}}$

***Similarity link prediction***

$Z^{RR}_{CLEAR} \leftarrow Z^{R}_{CLEAR} \,||\, Z^{R}_{CLEAR} \in \mathbb{R}^{2*\mathrm{h}}$

$Z^{DD}_{CLEAR} \leftarrow Z^{D}_{CLEAR} \,||\, Z^{D}_{CLEAR} \in \mathbb{R}^{2*\mathrm{h}}$

$Z^{TT}_{CLEAR} \leftarrow Z^{T}_{CLEAR} \,||\, Z^{T}_{CLEAR} \in \mathbb{R}^{2*\mathrm{h}}$, where $Z^{R}_{CLEAR}$, $Z^{D}_{CLEAR}$and $Z^{T}_{CLEAR}$ are CLEAR embedding of drug, disease and protein respectively

***Multi-task link prediction***

| | |
|---|---|
| Drug-Disease association | $(\hat{y}_{drug-disease}) \leftarrow MLP\ (Z^{RD}_{CLEAR})$ |
| Drug-Protein interaction | $(\hat{y}_{drug-protein}) \leftarrow MLP\ (Z^{RT}_{CLEAR})$ |
| Disease- Protein association | $(\hat{y}_{disease-protein}) \leftarrow MLP\ (Z^{DT}_{CLEAR})$ |
| Drug- Drug similarity | $(\hat{y}_{drug-drug}) \leftarrow MLP\ (Z^{RR}_{CLEAR})$ |
| Disease - Disease similarity | $(\hat{y}_{disease-disease}) \leftarrow MLP\ (Z^{DD}_{CLEAR})$ |
| Protein - Protein similarity | $(\hat{y}_{protein-protein}) \leftarrow MLP\ (Z^{TT}_{CLEAR})$ |
